# Adaptive Responses Directed by CREB Control Epithelial-Mesenchymal Plasticity in Cancer

**DOI:** 10.64898/2025.12.09.693241

**Authors:** Kshitij Parag-Sharma, Harish Bharambe, John J. Powers, Weida Gong, Daniel J. Purcell, Caleb Mensah, Chloe Twomey, Jay H. Mehta, Samy Lamouille, Antonio L. Amelio

## Abstract

Cellular plasticity plays essential roles in development including organogenesis and tissue homeostasis. The epithelial-to-mesenchymal transition (EMT) is no longer considered a binary switch but rather a dynamic process characterized by a continuum of metastable intermediates having unique features. This epithelial-mesenchymal (E/M) plasticity can be co-opted by cancer cells to promote dedifferentiation that results in hybrid E/M states which increase tumor heterogeneity and generate distinct molecular and phenotypic adaptations that promote drug resistance, dormancy, recurrence, and/or cell invasion and metastasis. The mechanisms that coordinate and maintain metastable hybrid E/M states are poorly understood, and here we report they are controlled by the master transcription factor CREB which regulates adaptive response genes necessary for E/M plasticity. Specifically, a CREB-dependent head and neck cancer model validated the role of CREB in cancer cell plasticity and revealed that it controls a non-canonical EMT gene signature. Moreover, analyses of this signature across cancer types identified the transcriptional regulators VGLL3 and KLF3 as core PanCancer mediators of hybrid E/M states, and gain- and loss-of-function studies established that CREB regulates E/M plasticity by coordinating VGLL3 and KLF3 to drive metastasis.

## Introduction

The epithelial-to-mesenchymal transition (EMT) is a dynamic process, where epithelial cells are reprogrammed to acquire mesenchymal phenotypes that imbues these cells with migratory abilities^1^. This phenotypic reprogramming occurs in response to context-dependent signals from the surrounding microenvironment and enables essential developmental processes to proceed during embryogenesis^2,3^. However, EMT is also associated with several diseases including cancer where these phenotypic changes facilitate tumor progression, metastasis, and resistance to therapeutic interventions^4,5^. While EMT was conventionally viewed as a bidirectional process yielding one of these two phenotypic states, accumulating evidence prompted the EMT International Association (TEMTIA) to issue revised guidelines regarding the existence of hybrid E/M states, sometimes referred to as partial or metastable EMT states^6^. This cellular plasticity in cancer has been coined epithelial-mesenchymal (E/M) plasticity and examples of this phenomenon can be traced back to early observations in breast carcinomas where neoplastic breast epithelial cells display variable differentiation trajectories, and by studies in breast and head and neck cancers showing that E/M plasticity contributes to cancer cell stemness, intra-tumoral heterogeneity, and metastatic potential^6–10^. However, the mechanisms that govern E/M plasticity and hybrid E/M states remain poorly defined.

## Results

### CREB transcriptional networks link adaptive signaling to E/M plasticity

Hormetic stressors that shape cellular plasticity via adaptive responses regulate many genes controlled by the master transcription factor cAMP Response Element Binding protein (CREB1) and its paralog Activating Transcription Factor 1 (ATF1)^11–15^. To assess if these adaptive cellular responses also govern E/M plasticity in cancer, canonical EMT genesets derived from the Molecular Signatures Database (MSigDB; Hallmark_EMT) and EMT signatures curated from two publicly available RNA-seq datasets (PanCancer EMT^5^ or biological inducers of context-specific EMT^16^) were used to investigate transcription factor enrichment at these core EMT genes. ChEA3 analysis of transcription factors associated with genes within these signatures revealed that CREB/ATF1 and several related family members are enriched across each EMT geneset, with 55 – 63% of EMT genes showing enrichment for 2 – 6 of these transcription factors (Fig. 1a, b, Extended Data Fig. 1). To test if EMT involves CREB, we established a genetic model in CREB-dependent salivary mucoepidermoid carcinoma (MEC) cells whereby conditional expression of the dominant negative molecule A-CREB^17^ is used to modulate the activity of endogenous CREB and its related bZip domain family members (Extended Data Fig. 2a-d). These MEC cells express an oncogene fusion (CRTC1-MAML2) that binds directly to the CREB bZip domain to activate genes bound by CREB. Consequently, inducible expression of A-CREB sequesters CREB and prevents transcriptional activation as confirmed by downregulation of the proto-typical CREB target gene, *NR4A2* (Extended Data Fig. 2e). Gene expression profiles were examined using bulk RNA-seq data generated from two independent cell lines with induced A-CREB expression at both acute and chronic time points (Fig. 1c, d). Gene set enrichment analysis (GSEA) of differentially expressed genes identified significant enrichment of the Hallmark_EMT pathway as well as hallmark pathways related to TNFα and TGFβ signaling which are known extrinsic regulators of EMT (Fig. 1e, Extended Table 1). To examine the effects of A-CREB on EMT gene expression, upregulated DEGs overlapping between the two cell lines were curated into a gene signature and the degree of overlap with canonical upregulated EMT genes was determined (Fig. 1f, g, Extended Tables 2, 3). Notably, a significant number of these genes do not overlap classical EMT genes; thus, non-canonical EMT-related genes are associated with CREB activity (Fig. 1h, Extended Table 2).

**Figure 1.**
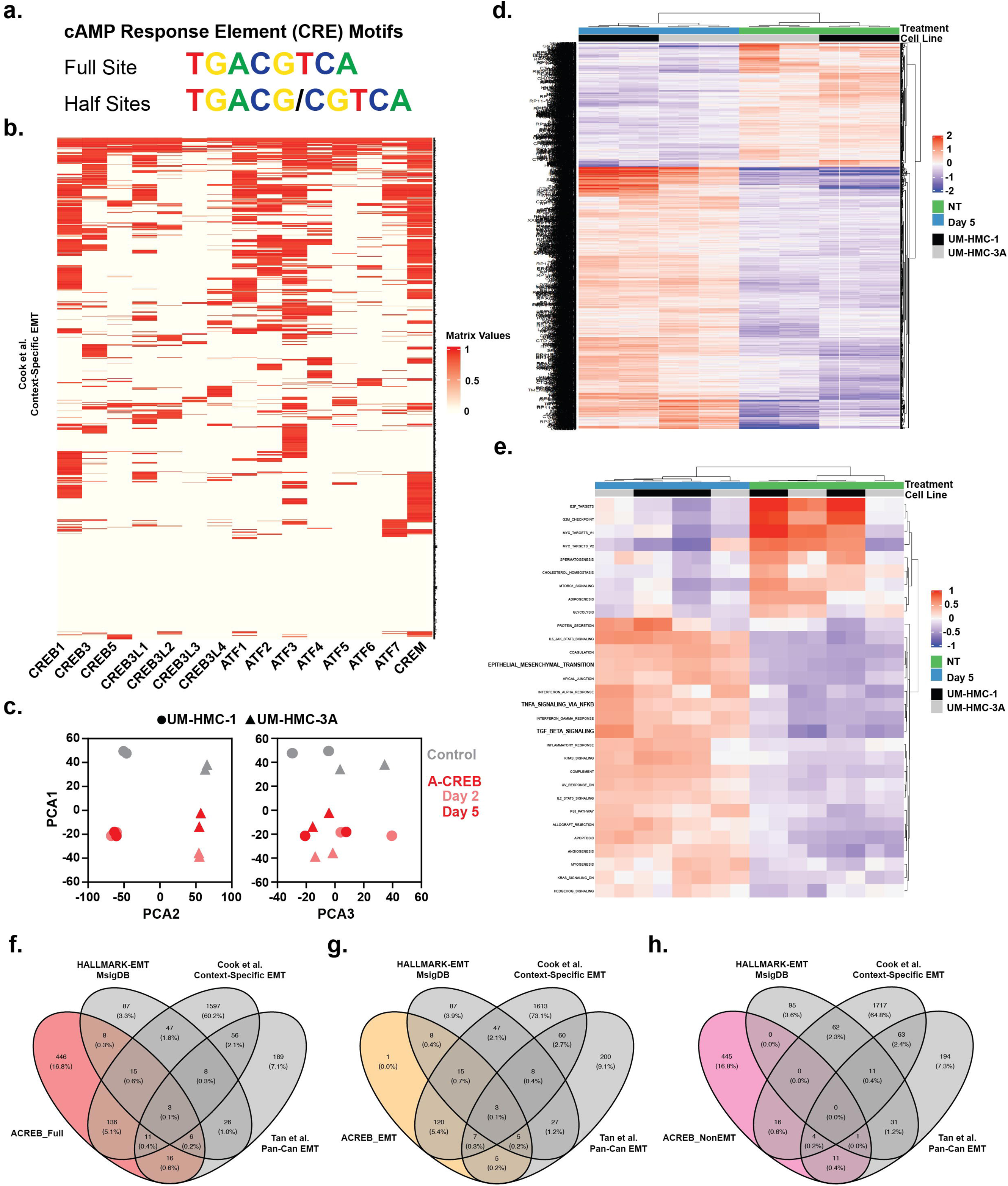
Identification of non-canonical EMT-related genes regulated by CREB. a, Representative full palindrome and half-site cAMP Response Element (CRE) motif sequences recognized by the CREB/ATF1 family of transcription factors. **b,** ChEA3 enrichment analysis of CREB/ATF1 family transcription factors. The interactive cluster-gram depicts enrichment of each transcription factor for differentially expressed genes related to Cook et al. context-specific EMT^16^. **c,** Principal component analysis of bulk RNA-sequenced UM-HMC cell lines annotated by control (NT) or Day 5 timepoint following doxycycline (Dox)-mediated induction of the dominant negative CREB molecule, A-CREB. **d,** Heatmap of DEGs between cell lines Dox-induced to express A-CREB and untreated control cells. Unsupervised hierarchical clustering performed on 2,961 DEGs with fold change > 1.5, padj < 0.01, and base mean > 10 (*n* = 2). **e,** MSigDB hallmark GSEA analysis of the UM-HMC cell lines expressing A-CREB at Day 5 versus control. The heatmap displays the enrichment scores for the 16 different hallmark gene sets with adj p-value < 0.01. **f-h,** Venn diagrams highlighting the overlap between upregulated canonical EMT genes and upregulated DEGs within the (**f**) ACREB_Full gene list, and the derivation of gene lists corresponding to canonical (**g**) ACREB_EMT and non-canonical (**h**) ACREB_NoEMT sub-signatures.

### Suppressing CREB drives EMT and metastatic potential

CREB signaling drives epithelial cell growth and proliferation and blocking CREB suppresses these activities and promotes differentiation^18,19^. However, roles for CREB/ATF1 have also been reported in upregulating mesenchymal genes such as fibronectin (*FN*) and slug/snail (*SNAI1* and *SNAI2*) that are important for cell migration and invasion^20–22^. To further examine the effects of blocking CREB activity on induction of EMT-related transcriptional programs, we induced A- CREB expression and monitored effects on proliferation, survival, and morphologic features of MEC cells. Blocking CREB impaired cell proliferation without affecting cell viability despite the presumed oncogene dependency on CREB signaling (Extended Fig. 3a, b). However, examination of cell morphology in two independent MEC cell lines revealed that blocking CREB leads to acquisition of classical EMT phenotypes characterized by the appearance of proto- typical mesenchymal cell morphology with front-back polarity, ruffled lamellipodia, and a marked decreases in direct cell-cell contact, which is in stark contrast to control cells which grow in colonies and display a cobblestone-like epithelial cell appearance (Fig. 2a). Moreover, A-CREB- expressing cells displayed a functional switch, with significant (> 4-fold) increases in single-cell motility and migratory speeds on the order of 2.35 ± 0.66 µm/min (Fig. 2b, c, Extended Videos 1, 2), which are comparable to those observed for some of the most motile cells recorded *in vitro*^23^. Furthermore, scratch assays demonstrated that these cells display ∼3x faster collective migration (Fig. 2d), and this is also associated with a dramatic increase in 2D and 3D invasive potential (Fig. 2e, f, Extended Fig. 3c).

**Figure 2.**
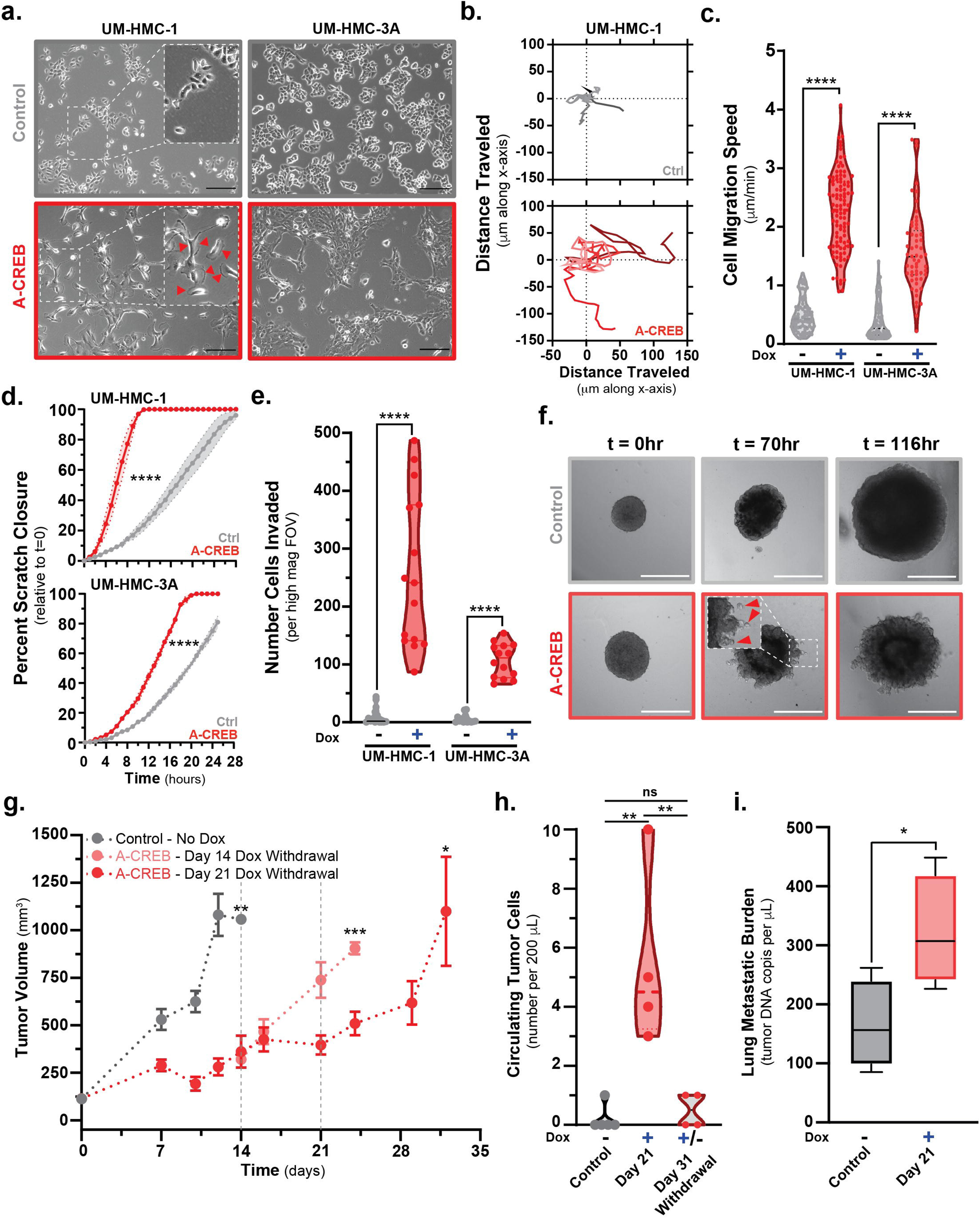
CREB disruption promotes EMT and metastatic behavior *in vivo*. a, Effects of disrupting CREB on cell morphology observed at 96 hours post-Dox addition to induce A-CREB versus control non-treated UM-HMC-1 and UM-HMC-3A cell lines. Red arrows within the high-magnification inset highlight the appearance of front-back polarity marked by ruffled lamellipodia (*n* = 3). **b,** Representative single-cell motility tracks of five randomly selected UM-HMC-1 cells that were either untreated controls or Dox-treated to induce A-CREB for 72 hours prior to tracking. **c,** Quantification of single-cell motility for UM-HMC-1 and UM-HMC-3A cell lines with or without A-CREB expression (*n* = 3, >50 cells analyzed per treatment condition, mean ± SEM, ** *p* < 0.0001). **d,** Time-lapse quantification of collective cell migration/scratch closure for UM- HMC-1 and UM-HMC-3A cell lines with or without induction of A-CREB expression (72 hours; *n* = 3, mean ± SEM plotted within error bars as shaded region, **** *p* < 0.0001). **e,** Quantification of transwell invasion assays for UM-HMC-1 and UM-HMC-3A cell lines with or without induced A-CREB expression (72 hours). Chambers with 8 μm pores were coated with 5% Matrigel prior to cell seeding (*n* = 3, mean ± SEM, **** *p* < 0.0001). **f,** Representative time-lapse images of 3D invasion outgrowth from tumor spheres embedded in Matrigel. Tumor spheres were established from UM-HMC-1 cells with or without A-CREB expression. Red arrows within the high-magnification inset highlight invading tumor cells (*n* = 3). **g,** Quantification of UM-HMC-1 orthotopic xenograft tumor growth in animals administered either normal chow for controls or Dox chow to induce A-CREB expression. Animals administered Dox chow were split into two groups with one group being switched back to normal chow after 21 days (*n* = 5, mean ± SEM, ** *p* < 0.01, *** *p* < 0.001, * *p* < 0.05). **h,** Quantification of GFP^+^TdTomato^+^ circulating tumor cells isolated from peripheral blood of orthotopically transplanted animals at the time points indicated (*n* = 4-5, mean ± SEM, ** *p* < 0.01). **i,** Quantification of metastatic tumor burden in the lungs of orthotopically transplanted animals at the time points indicated (*n* = 3, mean ± SEM, * *p* < 0.05).

Advanced salivary MEC commonly spreads to the loco-regional lymph nodes, yet the most common site for distant metastases is the lungs^24^. To test if the EMT phenotype induced upon blocking CREB is associated with increased metastatic potential, we developed an orthotopic salivary gland transplant model and observed that A-CREB induction reduces tumor volume at day 14 and day 21 post-Tx (Fig. 2g, Extended Fig. 3d). However, examination of peripheral blood at day 21 revealed significant increases in circulating tumor cells (CTCs), and switching animals back to normal chow at either day 14 or day 21 post-Tx led to rapid re-growth of the primary orthotopic tumor and to a decrease in CTCs (Fig. 2g, h, Extended Fig. 3e). Notably, while metastatic lesions were not detectable while animals were under Dox chow administration, analysis of resected lung tissues following Dox chow withdrawal revealed significant increases in distant lung metastatic burden (Fig. 2i). Thus, CREB/ATF1 controls successful colonization and growth at distant sites.

### CREB activity regulates hybrid E/M states and invasive potential

Plasticity of EMT states is associated with metastatic potential that is linked to distinct modes of 3D tumor cell migration involving hybrid E/M cells which retain partial epithelial phenotypes that invade and colonize as cell clusters^8,25–27^. Notably, examination of 3D tumor spheroid outgrowth at several time points revealed that salivary MEC cells with CREB disruption invaded in a collective fashion, as budding multicellular clusters rather than as spindle-like protrusions (Fig. 3a), suggesting that clusters retain some epithelial features (e.g., cell-cell contact) that are better suited to spread and colonize distant sites. To assess the extent to which CREB regulates the epithelial state and this mode of invasion, CREB activity was examined across independent MEC cell lines (Fig. 3b, c). These studies revealed that UM-HMC-1 cells having the highest CREB activity display ∼80-fold increased 2D spheroid spreading compared to UM-HMC- 3A cells that have lower basal CREB activity and are more effectively blocked by A-CREB (Fig. 3d, e, f).

**Figure 3.**
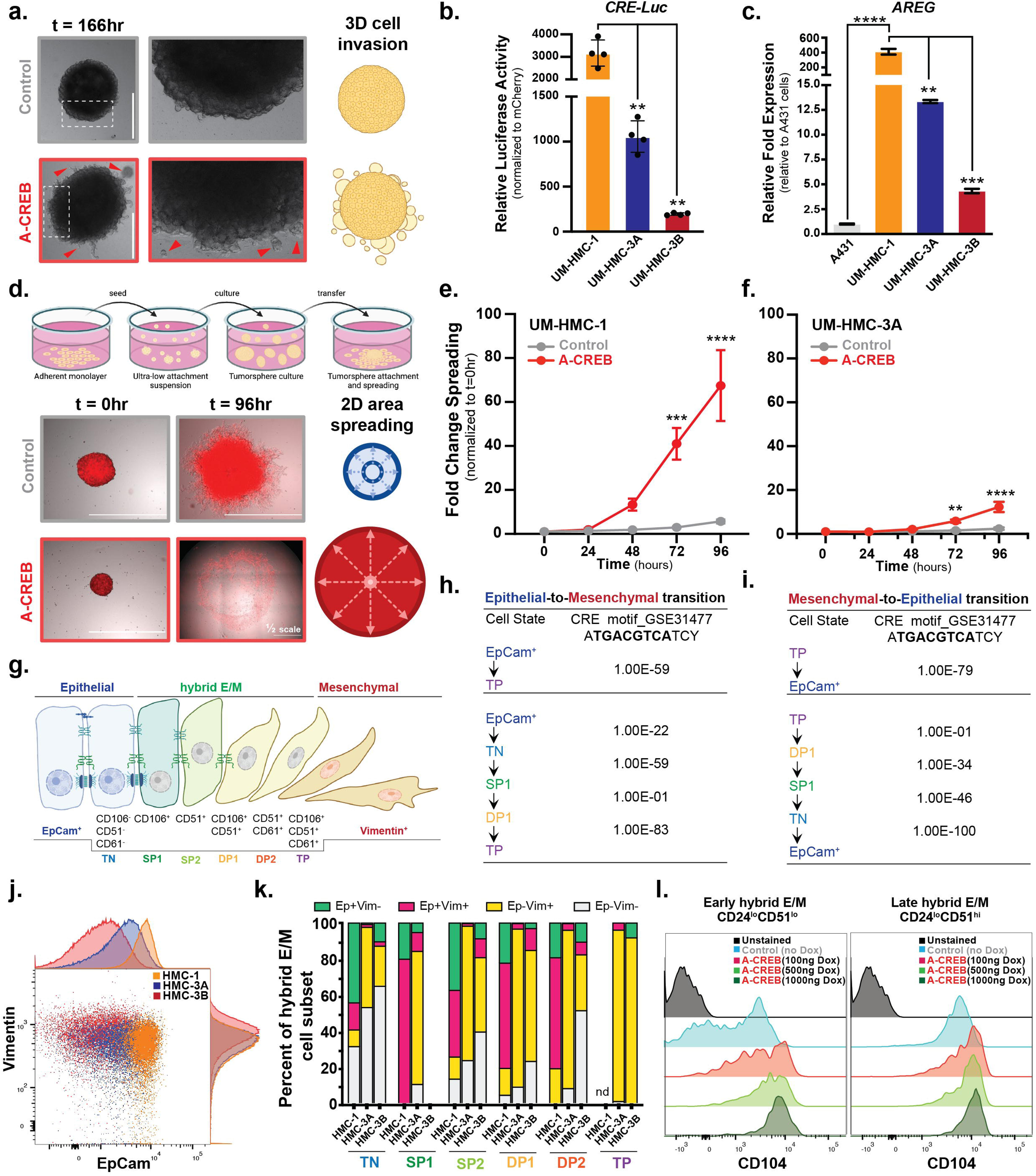
Hybrid Epithelial-Mesenchymal transition states are controlled by levels of CREB activity. a, Representative images highlighting collective 3D invasion of budding multicellular clusters from tumor spheres embedded in Matrigel. Tumor spheres were established from UM-HMC-1 cells with or without A-CREB expression. Red arrows within the high-magnification inset highlight invading tumor cell clusters (*n* = 3). **b-c,** Comparison of relative CREB activity within distinct UM-HMC cell lines. **b,** Transient co-transfections of each cell line were performed with the indicated cyclic AMP response element (CRE) luciferase reporter, and luciferase activity measured 48 h post transfection (*n* = 3, mean ± SEM, ** *p* < 0.01). **c,** Real-time qPCR of the indicated endogenous CREB target gene *AREG* in each cell line. Expression is shown normalized to *RPL23* and relative to the C1/M2 negative cell line A431 (*n* = 3, mean ± SEM, ** *p* < 0.01, **** *p* < 0.0001, *** *p* < 0.001). **d,** Schematic of 3D tumor spheroid spreading assay with representative images of UM-HMC-1 cells with or without A-CREB expression. **e-f,** Time-lapse quantification of 3D tumor spheroid spreading in (**e**) UM- HMC-1 cells with high basal CREB activity or (**f**) UM-HMC-3A cells with moderate basal CREB activity (*n* = 3, mean ± SEM, ** *p* < 0.01, *** *p* < 0.001, **** *p* < 0.0001). **g,** Diagram depicting the expression of cell surface markers present on epithelial, hybrid E/M, and mesenchymal tumor cells. Hybrid E/M cells express a combination of CD106 and/or CD51 in early hybrid E/M, or CD106, CD51, CD61, and/or CD104 in late hybrid E/M, while epithelial cells are triple negative (TN) for these markers and express EpCam, and late stage mesenchymal cells with full EMT phenotype are triple positive (TP) for these markers and also express Vimentin. **h-i,** Canonical CREB transcription factor motif enrichment in ATAC-seq datasets representative of EMT transition states as determined by Homer analysis by Pastushenko et al. The enrichment of CRE motifs was determined for peaks that were upregulated between the indicated subpopulations in cells undergoing (**h**) EMT or (**i**) MET. **j,** Marginal flow plots showing the basal EpCam and Vimentin expression levels according to CREB activity levels for the UM-HMC-1 (CREB^hi^), UM-HMC-3A (CREB^mid^), and UM-HMC-3B (CREB^lo^). **k,** Distribution of cells within each of the different EMT transition states represented as the proportion of cells expressing EpCam (Ep+/-) and/or Vimentin (Vim+/-), *n* = 3 biological replicates (nd = not detected). **l,** Flow cytometry profiles for UM-HMC-1 cells with or without A-CREB expression 72 hours prior to analysis of the proportions of hybrid E/M phenotypes. The EMT transition states were separated into early versus late hybrid E/M on the basis of CD24, CD51, and CD104 expression, *n* = 3 biological replicates.

The identification of distinct hybrid E/M transition states that are associated with the expression of various cell surface markers presented an opportunity to evaluate the contribution of CREB/ATF1 signaling in regulating E/M plasticity (Fig. 3g). Homer analysis of publicly available ATAC-seq data corresponding to these different hybrid E/M states revealed a significant enrichment of CREB motifs in peaks that were upregulated for all the indicated subpopulations undergoing EMT or mesenchymal epithelial transition (MET) (Fig. 3h, i, Extended Figs. 4a, b). Further, flow cytometry analysis of these populations revealed that CREB^hi^ UM-HMC-1 cells possess the highest proportion of cells expressing both epithelial (EpCam) and mesenchymal (Vimentin) markers, while this hybrid subpopulation is poorly represented in the UM-HMC-3A and UM-HMC-3B cells which have much lower CREB activity (Fig. 3j, Extended Figs. 4c, d, 5a).

To further define the relationship between CREB activity levels and the different hybrid E/M transition states, the various subpopulations were analyzed by flow cytometry (Extended Fig. 5b-g). Notably, the dual Ep^+^Vim^+^ subpopulation associated with CREB^hi^ UM-HMC-1 cells also display the highest proportion of hybrid E/M transition states, namely cells expressing the early single positive (SP1) CD106^+^ marker but also a significant proportion of the other SP2, DP1 and DP2 transition states (Fig. 3k). To confirm that CREB activity is required for these hybrid E/M transition states, varying doses of doxycycline were administered to titrate levels of CREB disruption in UM-HMC-1 cells engineered to inducibly express A-CREB. These studies confirmed that effectively blocking CREB with high levels of A-CREB led to a greater proportion of cells having a late CD24^lo^CD104^hi^CD51^hi^ hybrid E/M phenotype (Fig. 3l, Extended Fig. 5h-p), whereas low-level CREB inhibition enriched for cells having an early CD24^lo^CD104^lo^CD51^hi^ hybrid E/M phenotype. Thus, CREB activity directs E/M plasticity.

### Non-canonical PanCancer mediators of E/M plasticity are associated with CREB activity

To assess if there might be broader roles of CREB activity in E/M plasticity, and to identify effectors of CREB that control hybrid E/M transition states, a PanCancer analysis of additional cell lines corresponding to pancreatic adenocarcinoma, salivary squamous, head and neck squamous, mammary, epidermoid, and colorectal carcinomas was performed. These studies revealed that blocking CREB by inducible expression of A-CREB results in marked reduction in cell proliferation for all tumor cell lines tested, and in some cases these cells displayed phenotypic changes akin to those observed for the CREB-dependent UM-HMC cell lines (Extended Fig. 6a, b).

To investigate the prognostic potential of PanCancer gene signatures and their association with E/M plasticity, in-depth bioinformatics analyses were performed. These analyses revealed that a CREB-dependent non-canonical EMT gene signature (ACREB_NoEMT) connotes with poor prognosis across several cancers, similar to that observed with the Hallmark_EMT signature (Fig. 4a, b and Extended Fig. 6c). However, these four CREB-dependent cancers (glioblastoma, stomach, pancreatic, and kidney adenocarcinoma) represent some of the most aggressive and metastatic cancers, suggesting that the poor patient outcomes observed are merely a consequence of these cancers having high canonical EMTness. To address this, a list was curated of non-canonical ACREB-NoEMT genes shared between these four cancers and the CREB-related gene signatures (Fig. 4c, d, Extended Table 4). Notably, of the 477 upregulated genes in the non-canonical ACREB_NoEMT signature, there were six core overlapping shared genes (*KLK7*, *KRT16*, *MARCO*, *IL1RN*, *PTGS2*, and *VGLL3*). To examine the contribution of these six genes to EMT, we assessed their expression relative to the consensus Hallmark_EMT signature and found that only *VGLL*3 displayed a near complete PanCancer-wide (31 out of 33 TCGA cancers examined) positive correlation with tumor EMTness (Fig. 4e, Extended Fig. 7a).

**Figure 4.**
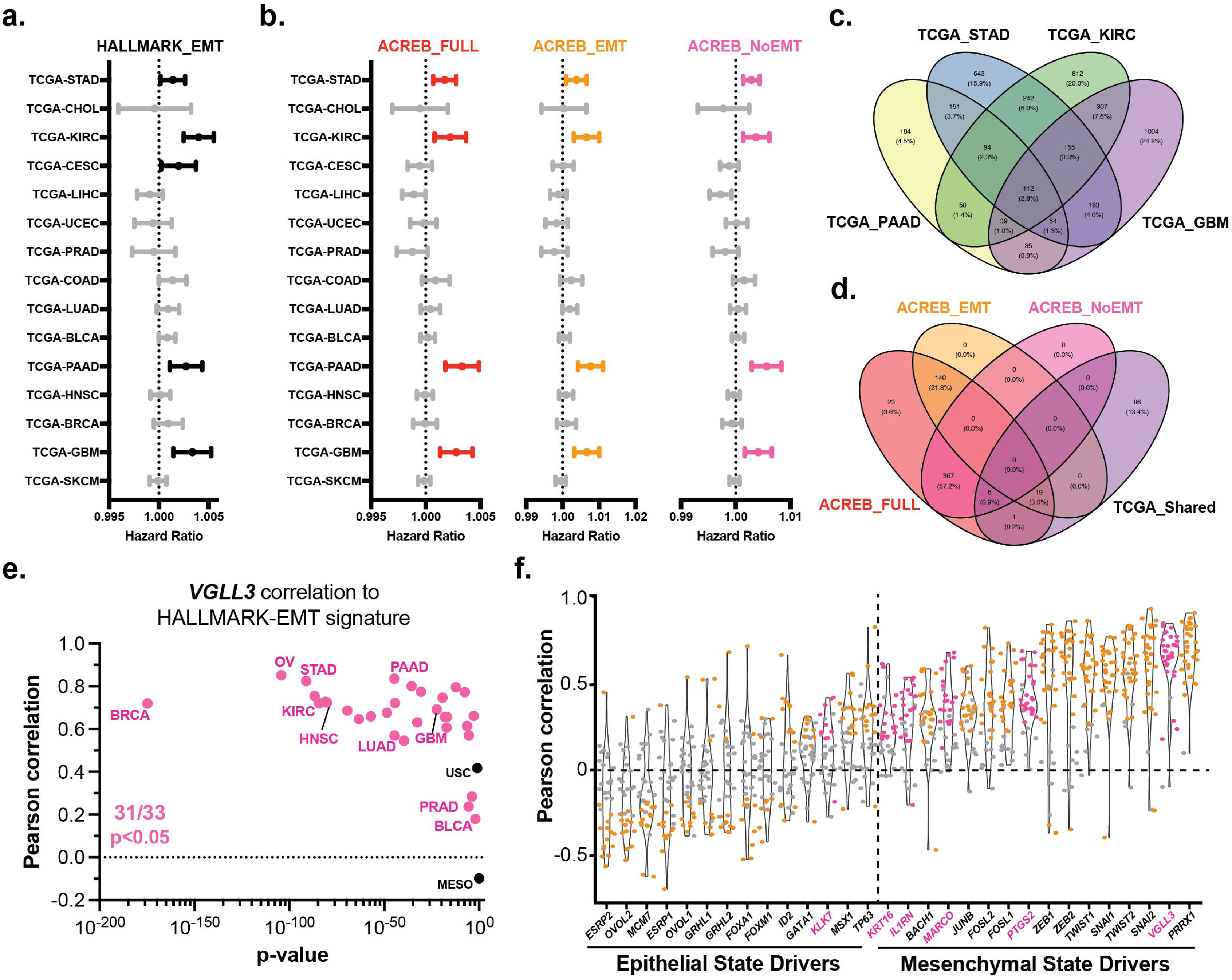
PanCancer identification of novel non-canonical regulators of EMT associated with CREB activity. a-b, Forest plots illustrating Hazard Ratios (HR) for the enrichment of genes associated with (**a**) Hallmark EMT or (**b**) the A-CREB_Full signature (red) versus signatures overlapping with canonical EMT genes for A-CREB_EMT (orange) or genes that do not overlap with canonical EMT for A-CREB_NoEMT (pink). Lines represent confidence intervals (95% CI) from multivariate Cox proportional hazard analysis for the TCGA datasets indicated with significant HR (p < 0.05) highlighted in bold. (**c**) Venn diagram used to identify 112 genes shared between the four TCGA cancers that display significant HR. (**d**) Venn diagram used to nominate 6 core non-canonical EMT genes shared between the shared TCGA cancer genes and the non-canonical A-CREB_NoEMT signature. (**e**) Correlation of gene expression for canonical regulators of epithelial versus mesenchymal cell states and each of the 6 core non-canonical A-CREB_NoEMT genes with the Hallmark_EMT gene signature. Pearson correlation plotted with respect to each canonical cell state regulator where each dot represents a unique cancer type across 33 TCGA cancers tested per cell state regulator. Grey dots are not significant, orange and pink dots are significant (p < 0.01) and reflect either canonical A- CREB_EMT genes or non-canonical A-CREB_NoEMT genes, respectively. (**f**) PanCancer correlation of gene expression for the shared non-canonical A-CREB_NoEMT core gene *VGLL3* with Hallmark_EMT gene signature. Pearson correlation plotted with respect to significance where each dot represents a unique cancer type across 33 TCGA cancers tested. Black dots are not significant and pink dots are significant (p < 0.05).

**Figure 5.**
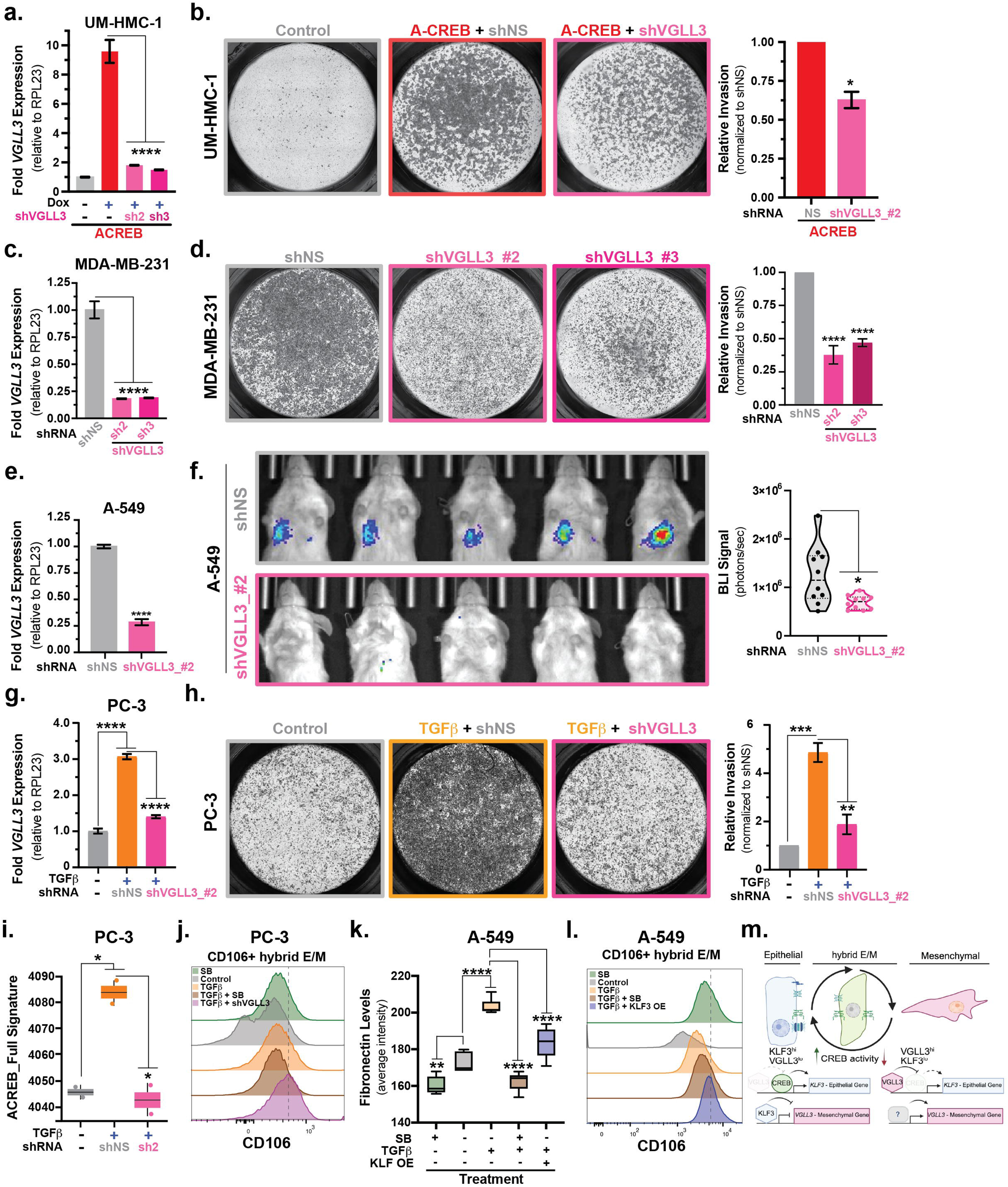
CREB activity coordinates *VGLL3* and *KLF3* expression to control hybrid E/M states that govern epithelial mesenchymal plasticity. a-b, shRNAs targeting *VGLL3* block the effects of A-CREB on inducing *VGLL3* expression and EMT-associated invasion. a, Real-time qPCR of endogenous *VGLL3* expression levels in stable UM-HMC-1 cells +/- Dox-induced A-CREB and +/- expression of two independent shRNAs. Fold expression is shown normalized to *RPL23* and relative to the no Dox control cells (*n* = 3, mean ± SEM, **** *p* < 0.0001). b, *Left*, representative transwell invasion assays with or without induced A-CREB expression and shRNA-targeted *VGLL3* knockdown 72 hours prior. *Right*, quantification of transwell invasion assays with or without induced A-CREB expression 72 hours prior. Chambers with 8 μm pores were coated with 5% Matrigel prior to cell seeding (*n* = 3, mean ± SEM, **** *p* < 0.0001). **c,** Evaluation of *VGLL3* knockdown on MDA-MB-231 breast cancer cells with high basal EMT-associated invasiveness. Two independent shRNAs targeting *VGLL3* were examined in stable knockdown cells to test the effects on gene expression by real-time qPCR of endogenous *VGLL3* expression levels. Fold expression is shown normalized to *RPL23* and relative to the no Dox control cells (*n* = 3, mean ± SEM, **** *p* < 0.0001). **d,** Representative images (*left*) and quantification (*right*) of transwell assays in MDA-MB-231 cells with stable *VGLL3* knockdown examined for effects on invasion. Chambers with 8 μm pores were coated with 5% Matrigel prior to cell seeding (*n* = 3, mean ± SEM, **** *p* < 0.0001). **e-f,** Effects of *VGLL3* knockdown on metastatic burden in the lungs of mice receiving tail-vein injections of A-549 cells stably expressing a LumiFluor bioluminescent reporter (GpNLuc) and either control non-specific (shNS) or *VGLL3*-targeting (shVGLL3_#2) shRNAs. **e,** Two independent shRNAs targeting *VGLL3* were examined by real-time qPCR in stable knockdown cells to assess effects on *VGLL3* expression levels. Fold expression is shown normalized to *RPL23* and relative to the control shNS cells (*n* = 3, mean ± SEM, **** *p* < 0.0001). **f,** Representative BLI images (*left*) and quantification (*right*) of metastatic burden in the lungs of mice receiving tail-vein injections of A- 549 cells stably expressing a LumiFluor bioluminescent reporter (GpNLuc) and either control non-specific (shNS) or *VGLL3*-targeting (shVGLL3_#2) shRNAs. BLI measurements were collected at endpoint from regions of interest and plotted (n = 5 per group, mean ± SEM, * *p* < 0.05). **g-h,** Experimental induction of EMT with TGFβ in PC-3 cells induces *VGLL3* gene expression and enhanced invasion which is reversible following knockdown. **g,** Real-time qPCR of *VGLL3* expression in untreated parental PC-3 cells or stable PC-3 cells expressing either control non-specific (shNS) or *VGLL3*-targeting (shVGLL3_#2) shRNAs treated with TGFβ stimulation. Fold expression is shown normalized to *RPL23* and relative to the untreated parental cells (*n* = 3, mean ± SEM, **** *p* < 0.0001). **h,** Representative images (*left*) and quantification (*right*) of transwell assays in PC-03 cells with stable *VGLL3* knockdown examined for effects on invasion. Chambers with 8 μm pores were coated with 5% Matrigel prior to cell seeding (*n* = 3, mean ± SEM, **** *p* < 0.0001). **i,** Flow cytometry profiles for control or *VGLL3* shRNA knockdown PC-3 cell lines treated with or without SB431542 or TGFβ for 72 hours prior to analysis of the CD106+ hybrid E/M phenotype (*n* = 2 biological replicates). **j,** ACREB_Full EMT signature enrichment scores in stable PC-3 cell lines expressing either control non-specific (shNS) or *VGLL3*-targeting (shVGLL3_#2) shRNAs (*n* = 2 per group). **k,** Untreated parental or stable A-549 cells overexpressing *KLF3* (KLF3 OE). Cells were treated with or without 5 ng/mL of TGFβ and/or 5 μM SB431542 (SB) for 72 h before fixation and immunofluorescence staining using an antibody against fibronectin. Cells were analyzed using an ImageXpress Pico Cell Imaging System, and the average fluorescence intensity in each well was determined using MetaXpress software (n = 6, ** *p* ≤ 0.01, **** *p* ≤ 0.0001). **l,** Flow cytometry profiles for untreated parental or stable A-549 cells overexpressing *KLF3* (KLF3 OE) and treated with or without SB431542 or TGFβ for 72 hours prior to analysis of the CD106+ hybrid E/M phenotype (*n* = 2 biological replicates). **m,** Model illustrating the role of CREB in controlling epithelial mesenchymal plasticity.

Given that EMT is a fluid process characterized by the dynamic interplay of transcriptional regulators controlling epithelial and mesenchymal states, we determined if the putative non-canonical drivers identified either 1) negatively regulate epithelial states (e.g., SNAI and ZEB) or 2) positively regulate mesenchymal states (e.g., TWIST and PRRX)^4^. Comparing the expression of these six genes to that of known regulators of epithelial versus mesenchymal cell states revealed that *VGLL3* displays minimal negative correlation to epithelial state genes but a strong positive correlation with mesenchymal state markers (Fig. 4f, Extended Fig. 7b, c). These data therefore suggest that VGLL3 is a positive regulator of mesenchymal states that is modulated by CREB activity.

### CREB orchestrates E/M plasticity via KLF3-mediated repression of *VGLL3*

Unlike the drosophila gene *vestigial* (*Vg*) that controls wing development, Vestigial-like 3 (*VGLL3*) is a mammalian ortholog that serves as a co-factor protein for TEA domain (TEAD) transcription factors that control a diverse array of pathways^28,29^. Importantly, recent studies demonstrated a role for VGLL3 in controlling cell proliferation and motility, and EMT-like phenotypes in cancer cells^30^. To test if *VGLL3* expression is necessary for driving the switch from an epithelial state to a more mesenchymal phenotype following attenuation of CREB signaling, we tested the effects of silencing *VGLL3* using two independent short hairpin RNAs (shRNA) and first evaluated their effects on EMT in UM-HMC-1 cells (Fig. 5a). *VGLL3* knockdown within UM-HMC-1 cells in combination with inducible A-CREB expression significantly reduced invasive potential compared to cells expressing the control non-specific shRNA (Fig. 5b). Notably, knockdown of *VGLL3* in eight tumor cell lines representing seven distinct cancer types having a more mesenchymal-like, invasive phenotype (breast, pancreatic, epidermoid, lung, head and neck, renal, and ovarian) demonstrated that *VGLL3* knockdown significantly impairs invasive behavior regardless of the tumor tissue-of-origin (Fig. 5c, d, Extended Fig. 8a-l), as well as metastatic potential *in vivo* (Fig. 5e, f, Extended Fig. 9a, b).

EMT is a non-linear, context-dependent process whereby E/M plasticity contributes to a high degree of variability among EMT programs^16^. One such program that is controlled by TGFβ signaling, provokes mesenchymal phenotypes in several systems^21,31,32^. Analysis of publicly available scRNA-seq data of A-549 lung adenocarcinoma cells demonstrated that *VGLL3* expression is induced by TGFβ treatment and drops to low levels following TGFβ withdrawal (Extended Fig. 9c). Potent induction of *VGLL3* by TGFβ was independently validated in PC-3 epithelial prostate cancer cells which display a dramatic EMT-mediated increase in cell invasion that is effectively blocked by shRNA-mediated knockdown of *VGLL3* (Fig. 5g, h). To validate these connections, bulk RNA-seq was performed on the TGFβ treated PC-3 cells with or without *VGLL3* knockdown, and in four tumor cell lines representative of distinct cancer types having mesenchymal-like phenotypes (MDA-MB-231, A-549, A-498, and T-24 cells). These analyses confirmed that silencing *VGLL3* is associated with significant decreases in CREB-mediated EMT genes (Fig. 5i, Extended Fig. 9d, e, Extended Table 5). Furthermore, *VGLL3* knockdown or antagonizing the effects of TGFβ-induced EMT by treatment with the potent inhibitor SB431542 led to increases in the early CD106+ hybrid E/M subpopulation (Fig. 5j, Extended Fig. 10a, b) that has been shown to promote distant metastases compared to other hybrid E/M subpopulations^8^.

Analysis of shared upregulated genes induced by *VGLL3* knockdown across these mesenchymal tumor cell lines identified the transcription factor Krüppel-like factor 3 (*KLF3*), which harbors clusters of binding motifs for CREB and the TEA domain family (TEAD) of transcription factors within the *KLF3* promoter (Extended Fig. 10c, d, Extended Tables 6, 7). KLF3 regulates epidermal differentiation by acting as a transcriptional repressor^33–37^, suggesting a feed-forward negative transcriptional circuit where VGLL3-TEAD represses, but CREB activates, *KLF3* expression which in turn represses *VGLL3* to maintain epithelial tumor cell states. Consistent with this model, prolonged A-CREB induction not only leads to elevated *VGLL3* levels but also reduced *KLF3* expression (Extended Table 1), and analysis of transcription factor binding sites identified by JASPAR revealed KLF3 binding motifs in the promoter proximal regulatory regions of *VGLL3* (Extended Table 8). To test this model, A549 cells were engineered to constitutively overexpress KLF3, and these cells showed significant decreases in cell invasion without affecting cell proliferation, consistent with these cells taking on a more epithelial-like phenotype (Extended Fig. 10e-h). Indeed, analysis of prototypical markers for epithelial (E-cadherin) and mesenchymal (N-cadherin) states revealed that KLF3 overexpression produced modest increases in E-cadherin levels and, more importantly, significantly impaired TGFβ-induced expression of N-cadherin (Extended Fig. 10i-k). Finally, KLF3 overexpression also significantly blocked TGFβ-induced expression of fibronectin and led to robust increases in the early CD106+ hybrid E/M subpopulation of TGFβ-treated cells (Fig. 5k, l, Extended Fig. 10l). Collectively, these data support a model whereby E/M plasticity is coordinated by levels of CREB activity which control hybrid E/M states via titration of *KLF3* and *VGLL3* expression (Fig. 5m).

## Discussion

CREB/ATF1 transcriptional networks regulate adaptive cellular responses, and here we demonstrate these include cancer-associated E/M plasticity. Specifically, inhibition of CREB signaling in MEC epithelial-like cancer cells impairs proliferation and induces morphological changes characteristic of EMT phenotypes, including marked increases in single-cell motility, invasion, and metastatic dissemination; thus, CREB maintains epithelial cell states. Importantly, modulating CREB activity revealed that it controls hybrid E/M states where retaining some CREB activity shifts cells toward distinct hybrid E/M cell phenotypes that express both epithelial and mesenchymal markers. Interestingly, PanCancer analyses unveiled non-canonical EMT genes such as *VGLL3* that are a hallmark of hybrid E/M states across broad tumor types, and functional studies revealed that CREB-mediated repression of *VGLL3* is necessary to sustain an epithelial cell state. Mechanistically, these data support a role for CREB in maintaining E/M plasticity by modulating expression of the transcriptional repressor *KLF3* which negatively regulates *VGLL3* expression. This CREB-dependent *KLF3*-*VGLL3* circuit controls hybrid E/M cell states involved in EMT transition linked to tumor cell migration, invasion, and metastasis. These findings position the CREB–KLF3–VGLL3 axis as a central regulator of cellular plasticity in cancer and suggest that targeted modulation of this pathway may provide novel strategies to limit metastasis and manipulate hybrid E/M dynamics therapeutically.

## Acknowledgments

The authors thank Drs. Robert Weinberg, John L. Cleveland, Andriy Marusyk, Jeremy Purvis, Richard Cheney, James Bear, and Colin O’Banion for helpful discussions during preparation of this manuscript, Gabriela De La Cruz and Bentley Midkiff in the Pathology Services Core, Alain Valdivia for technical veterinary assistance with the orthotopic salivary gland surgeries, Jennifer L. Modliszewski and David Corcoran in the Lineberger Bioinformatics Core of the University of North Carolina at Chapel Hill for expert technical assistance. The Pathology Services and Lineberger Bioinformatics cores are supported in part by an NCI Center Core Support Grant (P30-CA016086). This work was also supported in part by Jodi Kroger, Bethany Carter, and Neelkamal Chaudhary of the Flow Cytometry Core, Jodi Balasi of the Tissue Core, and Dr. Mikalai Budzevich and Epi Ruiz of the Small Animal Imaging Lab at the Moffitt Cancer Center and Research Institute, a comprehensive cancer center designated by the National Cancer Institute and funded in part by Moffitt’s Cancer Center Support Grant (P30-CA076292). This work was supported in part by a UNC Dissertation Completion Fellowship (to KPS), Moffitt Cancer Center funds (to ALA), and NIH/NIDCR R01-DE030123 (to ALA).

## Author contributions

K.P.S., H.S.B., and A.L.A. designed the experiments and performed data analysis. K.P.S. and H.S.B., performed most of the biological experiments and the *in vivo* animal studies. H.S.B. and J.J.P. performed flow cytometry and analyses for the hybrid E/M markers. K.P.S. and C.M.T. performed the ImageJ analyses and quantification. W.G. and J.H.M. performed the bioinformatic analysis. D.J.P., C.M., and S.L. performed the immunofluorescence and immunoblotting for EMT markers. K.P.S., H.S.B., and A.L.A. wrote the initial draft of the manuscript. All authors read, edited, and approved the final manuscript.

## Methods

### Animal studies

All animal studies were approved by the Institutional Animal Care and Use Committees (IACUC) of The University of North Carolina Chapel Hill, Moffitt Cancer Center, and the University of South Florida. Male 6-8 week old athymic nude mice (Nu/Nu) were obtained from the Animal Studies Core at the University of North Carolina at Chapel Hill and housed in facilities run by the Division of Comparative Medicine at the University of North Carolina (Chapel Hill, NC, USA). For all xenograft studies, mice were subcutaneously injected with 1x10^6^ UM- HMC cells resuspended in 50% HBSS and 50% Matrigel. Caliper measurements were collected every 2 days and bioluminescent imaging (BLI) was performed every 5 days throughout the course of the study. For all orthotopic salivary gland xenografts, 8-week old NSG immunocompromised animals were obtained from the Animal Studies Core at UNC-CH and housed in facilities run by the Division of Comparative Medicine core at UNC-CH. Mice were anesthetized using isoflurane and a 4 mm incision was made above the salivary glands (see Extended Fig. 3d). 25,000 HMC-1 MEC cells were injected in a total volume of 30 µL (15 µL cells in HBSS + 15 µL Matrigel) into the left submandibular gland, and the incision was sutured closed. Animals were given the analgesic meloxicam and PBS subcutaneously every day for three days to soothe post-surgery pain and maintain proper hydration, along with wet food for one-week post-surgery. Tumor growth was measured twice a week using calipers once tumors were palpable. As indicated, animals were switched to, and maintained on, Dox-laden chow once the average tumor sizes exceeded 100 mm^3^. At endpoints, before sacrifice, animals were bled sub-orbitally to collect blood for analysis of circulating tumor cells. Animals were sacrificed using CO_2_ euthanasia and cervical dislocation.

### Cell lines

UM-HMC-1, UM-HMC-3A, and UM-HMC-3B cells were kindly provided by Dr. Jacques Nör (University of Michigan, Ann Arbor, MI, USA)^38^. A-253, A-388, A-431, A-498, A- 549, BxPC-3, UM-SCC-5, MCF-7, HCT116, MDA-MB-231, PSN-1, T-24, 786-O, UM-SCC-74A, SK-OV-3, and PC-3 were acquired from the UNC-CH Lineberger Cancer Center Tissue Culture Facility (TCF) on-site cell line inventory. UM-HMC-1 (source: male, salivary gland mucoepidermoid carcinoma (MEC)), UM-HMC-3A (source: female, hard palate MEC), and UM- HMC-3B (source: female, lymph node metastasis of hard palate MEC) parental and stably transduced lines were cultured in DMEM medium (GIBCO #11965-118) supplemented with 10% FBS (Atlanta Biologicals #S11550), 20 ng/mL EGF (Sigma-Aldrich #E9644), 400 ng/mL hydrocortisone (Sigma-Aldrich #H0888), 5 μg/mL insulin (Sigma-Aldrich #I6634), and 1X pen/strep/glutamine (PSG; Life Tech #10378016). A-388 (source: male, squamous cell carcinoma (SCC)), A-498 (source: male, renal cell carcinoma), A-549 (source: male, lung adenocarcinoma), A-431 (source: female, skin SCC), UM-SCC-74A (source: male, tongue SCC), UM-SCC-5 (source: male, laryngeal SCC), HCT-116 (source: male, colon carcinoma), MDA-MB-231 (source: female, breast adenocarcinoma), MCF-7 (source: female, invasive breast carcinoma of no special type), SK-OV-3 (source: female, ovarian serous cystadenocarcinoma), and HaCaT (source: male, spontaneously immortalized keratinocyte) cells were cultured in DMEM supplemented with 10% FBS, 1X GlutaMAX, 1X HEPES (Corning #25-060-CI), and 1X PSG, with the exception of the SK-OV-3 which also received 1X non-essential amino acid (NEAA) supplementation. A-253 (source: male, salivary gland SCC) cells were cultured in McCoy’s 5A medium (Life Tech #16600108) supplemented with 10% FBS, 1X GlutaMAX, and 1X PSG. BxPC-3 (source: female, pancreatic ductal adenocarcinoma), PSN-1 (source: male, pancreatic adenocarcinoma), 786-O (source: male, renal cell carcinoma), T-24 (source: female, bladder carcinoma), and PC-3 (source: male, prostate carcinoma) cells were cultured in RPMI (GIBCO #11875093) supplemented with 10% FBS, 1X GlutaMAX, 1X HEPES, and 1X PSG. Cells were passaged using TrypLE (GIBCO #12604013) every 2-3 days or when they reached 90% confluence. All cells were maintained in a 37°C, 5% CO_2_ atmosphere. All cell lines were confirmed mycoplasma-free by PCR using mycoplasma detection primers as previously described^39,40^.

### Plasmids and cloning

The pRRL-rtTA3 lentiviral vector was kindly provided by J. Zuber, Research Institute of Molecular Pathology, Vienna, Austria. CMV500 A-CREB was a gift from Charles Vinson (Addgene plasmid #33371). pZeoSV2(+) vector backbone was purchased from Invitrogen (cat.#V85001). pCMV7.1-3xFLAG vector backbone was purchased from Sigma (cat.#E4026). The plasmid containing the ORF for HumanVGLL3-Full Length (isoform1, NM_016206.3) was purchased from GeneCopoeia (cat.#EX-Y3981-M02-B). The pTight:ACREB_PGK:TdTomato construct was generated as previously described^40^ by directionally cloning cDNA for the dominant negative CREB molecule A-CREB^17^ via 5′ *Not*I and 3′ *Mlu*I restriction enzyme sites into pRetroX-Tight-tdTomato^41^. The pRetroX_TRE3G:MCS_PGK:GpNLuc construct was generated by excising the pTight (rtTA Gen2) promoter from Addgene #70185 using XhoI (NEB Biosciences cat.#R0146) and BamHI (NEB Biosciences cat.#R0136), via sequential digestion. BamHI digested linearized plasmid was first blunted using T4 polymerase (NEB Biosciences cat.#R0203) prior to the XhoI digestion. The doubly digested backbone plasmid was treated with rSAP (NEB Biosciences cat.#R0371) and the vector was gel purified (Macherey-Nagel cat.#704609).

The TRE3G (rtTA Gen3) was excised from Addgene plasmid #85449 using XhoI and SmaI and gel purified (Macherey-Nagel cat.#704609). The TRE3G promoter was ligated into the digested/linearized/rSAP treated MCS_PGK:GpNluc vector using T4 DNA ligase (NEB Biosciences cat.#R0202). Ligation was transformed into home-made competent (Zymo Research cat.#M3015-500) E.coli Stable3 cells (ThermoFisher Scientific #C737303) and seeded onto Ampicillin (Corning cat.#46-100-RG) containing agar plates. Multiple colonies were picked, miniprepped (Macherey-Nagel cat.#740588) and sequence verified (Eton Biosciences Inc.). All enzymatic reactions, gel purifications and plasmid extractions were performed following the manufacturers recommended protocols.

Plasmid constructs expressing shRNAs targeting VGLL3 were obtained from the MISSION shRNA library (Sigma-Aldrich): shVGLL3_#1 (TRCN0000146645; sequence: CTTTCATGGAACAGTAGACAT) and shVGLL3_#2 (TRCN0000147610; sequence: GAATAGTTTCCCAACTTCCTT).

The KLF3 overexpression construct was obtained was kind gift from Dr. Prashant Mali (Addgene plasmid #120494). To generate a C-terminally tagged construct, a 1X FLAG epitope was introduced immediately upstream of the stop codon using an end-to-end amplification strategy. The FLAG sequence was incorporated into both forward and reverse primers (Forward: gatgatgataaaGGATCCGGCGCAACAAAC; Reverse: atctttataatcGACTAGCATGTGGCGTTTC). The entire parental plasmid was amplified with these primers using Q5 High-Fidelity DNA polymerase (New England Biolabs, M0491S). The resulting PCR product was purified, end-phosphorylated with T4 polynucleotide kinase, self-ligated with T4 DNA ligase, and transformed into ultracompetent *E. coli* Stbl3 cells. Positive clones were selected with 100 µg/ml ampicillin, and insertion of the FLAG tag along with overall sequence integrity was verified by whole-plasmid sequencing (Genewiz).

### Virus packaging, transduction, and stable cell line generation

Virus packaging and cell transduction was carried out as previously described^40^. Briefly, the required lentiviral or retroviral plasmids were co-transfected with VSV-G envelope plasmid and either δ8.2 gag/pol (lentivirus) or pMD (retrovirus) helper plasmids into Lenti X-293T cells (Takara cat.#; 632180) seeded in 0.1% gelatin coated 10 cm tissue culture dishes (at ∼75% confluency). Transfections were performed using 1 mg/mL Polyethyleneimine (PEI) Transfection Reagent (VWR #BT129700) as previously described. Briefly, 1.5 μg VSV-G, 5 μg δ8.2 or pMD, and 6 μg lentiviral or retroviral plasmid were brought added to 500 μL of OptiMEM (LifeTech #1158021) and vortexed briefly. In a separate tube, 25 μL PEI (2 μL PEI/μg DNA) was added to 475 μL OptiMEM and vortexed briefly. Both solutions were incubated at room temperature for 5 min, combined and incubated for an additional 20 min at room temperature. PEI+DNA mixture was then added dropwise to the seeded Lenti X-293T cells. The next day, cell culture media was replaced with DMEM supplemented with 1x NaPyr, 10 mM HEPES, 1X GlutaMAX, and 1X PSG (no FBS). Two days later, media was collected and filtered through a 0.45 μm PVDF membrane and viral particles were concentrated via ultracentrifugation at 100,000 g for 2 hr at 4°C, into a sucrose cushion. Concentrated virus was resuspended in cold OptiMEM and either stored at −80°C or used immediately for transduction. ∼1/4 to 1/6^th^ of a p10 dishes worth of virus was used to transduce cell lines as needed to obtain a ∼60-80% transduction efficiency.

Alternatively, virus was packaged in 6-well plates using lipofectamine3000 (Invitrogen cat.# L3000015). 0.3 μg VSV-G, 1 μg δ8.2, and 1.2 μg lentiviral plasmid + 7.5 µL of p3000 Enhancer reagent were added in 125 uL of OptiMEM and vortexed briefly. In a separate tube, 7.5 μL lipofectamine 3000 (3 μL lipo3000/μg DNA) was added to 125 μL OptiMEM and vortexed briefly. Both solutions were incubated at room temperature for 5 min, combined and incubated for an additional 20 min at room temperature. Lipo3000+DNA mixture was then added dropwise to the seeded Lenti X-293T cells. The next day, cell culture media was replaced with DMEM supplemented with 1x NaPyr, 10 mM HEPES, 1X GlutaMAX, and 1X PSG (no FBS). Two days later, media was collected and filtered through a 0.45 μm PVDF membrane. Virus particle-containing media was either used fresh or frozen down at -80C for later use. Virus was used at 25-50% to transduce cell lines as needed to obtain a ∼60-80% transduction efficiency.

To generate stable cells, cells were seeded at 50-70% confluency in 6-well plate. Virus particles and 4 μg/mL polybrene (final) were added to ∼3-4 mL of media, mixed by pipetting and added to cells. Plates were centrifuged at 1200 g for 90 min at 30°C. 48 hours post transduction cells were split using TrypLE and expanded for downstream use (antibiotic selection or FACS sorting). Stable cell lines were validated by confirming doxycycline mediated protein induction by treating cells with 1 µg/mL doxycycline for 48-72 hours followed by Western blotting using the appropriate antibodies, or by qPCR to validate on-target knockdowns. Cells were selected using puromycin (Gibco Invitrogen, cat.#A11138-03) for >7 days, UM-HMC-1 (0.4 µg/mL), UM-HMC- 3A (0.4 µg/mL), A-549 (3 µg/mL), UM-SCC-74A (1 µg/mL), MDA-MD-231 (1 µg/mL), T-24 (0.5 µg/mL), 786-O (1 µg/mL), A498 (1 µg/mL), SK-OV-3 (1 µg/mL), PSN-1 (0.6 µg/mL).

### Real-time qPCR

mRNA was extracted from cells using the Nucleospin RNA mini kit (Macherey-Nagel, cat.# 740955). 0.5-2 µg of mRNA was converted into cDNA (BioRad cat.#1708891). For each reaction, 20 ng of cDNA (2 µL) was used per well for qPCR in a total reaction volume of 10 µL. qPCR was performed using KAPA SYBR Fast Master Mix (Roche, cat.# KK4610) in a C1000 thermocycler (BioRad). Relative gene expression of target genes was quantified using the 2ΔΔCt method, human RPL23 expression was used as a housekeeping gene to normalize target gene expression levels. All enzymatic reactions and sample extractions were performed following the manufacturer’s recommended protocols.

### Cell proliferation assay

Cells were seeded in a 6 well plate (UMHMC-1= 125,000 cells/well and UMHMC-3A= 250,000 cells/well) in media +/- doxycycline. Fresh media +/- doxycycline was added daily. 72 hours post seeding cells were trypsinized in TrypLE and total cell counts were quantified (BioRad TC20 automated cell counter) and graphed as Day 3/Day 0 cell count ratio.

### Collective cell migration assay

Doxycycline inducible A-CREB MEC cell lines were seeded in a 6 well plate (UM-HMC-1= 175,000/well or UM-HMC-3A= 350,000/well) in media with or without 1 µg/mL doxycycline. Cells were cultured for 3 days, with wells being washed and fresh media added daily. 72 hours post initial seeding/doxycycline addition, cells were trypsinized, counted and seeded for the scratch assay. Ibidi 2 well culture inserts (Ibidi, cat#; 81176) were placed in 24 well dish and cells (UM-HMC-1= 25,000/side or UM-HMC-3A= 40,000/side) were seeded in a 100 µL volume per side per chamber. 24 hours post seeding, once cells had formed a 100% confluent monolayer, ibidi culture inserts were gently and carefully removed from the wells. Wells were washed once with 1xDPBS and 1 mL of fresh media with/without doxycycline was added. Collective migration/scratch closing was imaged once every hour over a period of 24 hours using a Cytation 5 plate imager (BioTek). Wound size was quantified manually using the area measurement function in ImageJ software.

### 2D random cell migration

Doxycycline inducible A-CREB MEC cell lines were seeded in a 6 well plate (UM-HMC-1= 175,000/well or UM-HMC-3A= 350,000/well) in media with or without 1 µg/mL doxycycline. Cells were cultured for 3 days, with wells being washed and fresh media added daily. 72 hours post seeding, cells were trypsinized, counted and seeded for the random cell migration assay. Cells were seeded in a 24 well dish (UM-HMC-1= 2,500/well or UM-HMC- 3A= 5,000/well) in media with/without doxycycline. 24 hours post seeding, wells were washed once with 1xDPBS and fresh media with/without doxycycline added. Cells were imaged every 10 minutes over a period of 24 hours using a Cytation 5 plate imager (BioTek). Individual cell migration was tracked using the manual tracking plugin in ImageJ software. In order to calculate reliable and long-term average migration speeds, cell migration speed for an individual cell was calculated by averaging its speed across all timepoints.

### Transwell chemotaxis invasion assay

Doxycycline inducible A-CREB MEC cell lines were seeded in a 6 well plate (UM-HMC-1= 175,000/well or UM-HMC-3A= 350,000/well) in media with or without 1 µg/mL doxycycline. Cells were cultured for 3 days, with wells being washed and fresh media added daily. 72 hours post seeding, cells were trypsinized, counted, washed and resuspended in a low-serum media. Low-serum media was made by diluting the complete UM-HMC growth media 1:20 in DMEM (+ HEPES + GluaMAX + Sodium pyruvate, no FBS or additional growth factors). Prior to seeding cells onto the transwell chamber (Falcon permeable- transwell chambers- 8 um pore membrane, cat#; 353097), the membrane was first coated with Matrigel. Briefly, a 100 uL of 5 % Matrigel solution was added to the transwell chamber, which was then incubated for 1 hour in a humidified 37C, 5% CO_2_ cell culture incubator, to ensure polymerization. After polymerization the transwell chamber was placed into a 24 well plate well containing complete 750 µL of complete UM-HMC cell culture media (chemoattractant). Cells were seeded (UM-HMC-1= 100,000/transwell or UM-HMC-3A= 250,000/transwell) into the transwell chamber in 750 µL of low-serum media and allowed to invade through the Matrigel and onto the bottom side of the transwell membrane overnight. The next morning, fresh media was added to the wells and chambers were harvested ∼8-9 hours later by fixing cells using 10% formalin for 5 minutes at room temperature. Cells were visualized using 0.05% crystal, 30 minute incubation at room temperature followed by five washes using distilled water. Non-invaded cells still adhered to the ‘top-side’ of the transwell chamber were removed using cotton swabs. Membranes were allowed to dry overnight, and invaded cells were imaged in brightfield using a Cytation 5 plate imager, and relative cell invasion was quantified by measuring % of the chambers with invaded cells using ImageJ.

For VGLL3 knockdown experiments the following cell numbers were used per chamber, UM-HMC-1 (100,000/transwell), A-549 (125,000/transwell), UM-SCC-74A (50,000/transwell), MDA-MD-231 (125,000/transwell), T-24 (75,000/transwell), 786-O (50,000/transwell), A498 (125,000/transwell), SK-OV-3 (125,000/transwell), PSN-1 (125,000/transwell). For VGLL3 knockdown + TGFβ stimulated invasion assays (PC-3), 400,000 cells were seeded in a 6 well plate and treated with 5 ng/mL TGFβ (Sigma-Aldrich cat.#T7039) 24 hours post-seeding. 72 hours post TGFβ addition cells were trypsinized and seeded for invasion (250,000 cells/transwell) as described above. For TGFβ stimulated invasion assays (A-549), 200,000 cells were seeded in a 6 well plate and treated with 10 ng/mL TGFβ (Sigma-Aldrich cat.#T7039) 24 hours post-seeding. 72 hours post TGFβ addition cells were trypsinized and seeded for invasion assays as described above.

### 3D tumor spheroid spreading assay

Single MEC tumor spheroids were generated using the hanging drop culture method. Briefly, MEC cells (UM-HMC-1= 3,000/drop or UM-HMC-3A= 1,000/drop) were culture as a hanging drop in 30 µL of media. 3D tumor spheroids could form for 96 hours post seeding. A-CREB expression was induced by ‘spiking’ the cultures with doxycycline 24 hours post seeding, 10 µL of media with or without doxycycline was added to each drop, to a final concertation of 1 ug/mL doxycycline. Given the short half-life of doxycycline, an additional 10 µL ‘spike-in’ was added 72 hours post seeding (48 hours post the first doxycycline addition), to ensure continuous A-CREB induction. 96 hours post seeding, the 3D spheres were combined with 50 µL of Matrigel (∼1:1 dilution), and the 50% Matrigel + 3D sphere mix was seeded into a 96 well clear bottom opaque black plate. Spheroids were centrifugated briefly to enable contact between the spheroid and the bottom of the plate (∼500g for 1 minute). Plate was placed into a humidified, 37°C, 5% CO_2_ cell culture incubator for 1 hour to ensure Matrigel polymerization. After polymerization, 150 µL of full cell culture media was added per well, with/without doxycycline (1 mg/mL final). 3D spheroids were imaged every 4 hours. Spheroids were imaged as montages using the Cytation5 (BioTek) plate reader (brightfield and Texas Red filter cubes). 50% Matrigel was used to minimize 3D tumor spheroid invasion and favor 2D spreading across the well bottom.

### 3D tumor spheroid invasion assay

Single MEC tumor spheroids were generated using the hanging drop culture method. Briefly, MEC cells (UM-HMC-1= 3,000/drop or UM-HMC-3A= 1,000/drop) were culture as a hanging drop in 30 µL of media. 3D tumor spheroids could form for 96 hours post seeding. A-CREB expression was induced by ‘spiking’ the cultures with doxycycline 24 hours post seeding, 10 µL of media with or without doxycycline was added to each drop, to a final concertation of 1 mg/mL doxycycline. Given the short half-life of doxycycline, an additional 10 µL ‘spike-in’ was added 72 hours post seeding (48 hours post the first doxycycline addition), to ensure continuous A-CREB induction. Prior to transferring the spheroid into the 96-well plate, wells were coated with 100 µL of 2% agarose and allowed to polymerize at 4°C for 10 minutes. 96 hours post seeding, the 3D spheres were harvested in a total of 80 µL media and combined with combined with 20 µL of Matrigel (20% final Matrigel), following which the Matrigel + 3D sphere mix was seeded into a 96 well clear bottom opaque black plate. Plate was placed into a humidified, 37°C, 5% CO_2_ cell culture incubator for 1 hour to ensure Matrigel polymerization. After polymerization, 150 µL of full cell culture media was added per well, with/without doxycycline (1 mg/mL final). 3D spheroids were imaged every 4 hours. Spheroids were imaged as Z-stacks using the Cytation5 (BioTek) plate reader (brightfield and Texas Red filter cubes).

### Circulating tumor cell quantification

Blood was harvested from mice at the indicated time points either using retro orbital bleed or end point cardiac puncture (https://www.jove.com/v/10246/blood-withdrawal-i). >200 µL of blood was recovered for each mouse and rapidly transferred to a heparin coated collection tube. RBCs were lysed using ACK lysis buffer (Gibco, cat#A1049201) following manufacturers’ recommendation. Briefly, excess lysis buffer (1:10) was added to each sample and samples were incubated at room temperature for 5 minutes, with gentle vortexing every 30 seconds. Samples were spun down at 1000g for 5 minutes. Supernatant was aspirated carefully, pellets were resuspended in 10 mL of ice cold 1xHanks Buffered Saline Solution (Gibco, cat#14025092). Resuspended samples were spun down at 500g for 5 minutes and supernatants were carefully aspirated. Pellets were resuspended in 50 µL of 1xHBSS and resuspended samples were sandwiched between two slides (Fisher Scientific cat#; 1255015). Slides were manually line scanned on a Leica microscope using the BFP, GFP, and TdTomato channels. Circulating tumor cells were defined as GFP (GpNLuc) and TdTomato (A-CREB) positive and negative for BFP (autofluorescence) signal.

### Tumor tissue harvesting and fixation

Tumors were excised and chunks were flash frozen, on dry ice, for mRNA and protein lysates. The remaining tumor was cut into 4-8 pieces and fixed in 10% neutral buffered formalin for ∼1 week at room temperature. Tissues were harvested (lung, liver, lymph nodes and contralateral ‘uninjected’ salivary gland) and processed for *ex vivo* bioluminescent imaging. Following *ex vivo* BLI, all tissues were fixed in 10% neutral buffered formalin for approximately 1 week at room temperature. At the end of 1 week fixation, samples were washed in de-ionized water twice, for 2 days each, and incubated in 70% ethanol for 2 days to dehydrate them. Following fixation and dehydration, tissues were processed on an ASP6025 automated tissue processor (Leica Biosystems) and embedded in paraffin wax.

### Quantification of lung metastases by digital droplet PCR

Paraffin embedded whole lungs were extracted by heating the blocks to 95°C for 3 minutes. Paraffin embedded lungs were de- paraffinized by incubating in xylene (∼25 mL), briefly, lungs were incubated in xylene overnight at room temperature under constant rocking, the following day fresh xylene was added to the tubes and samples were incubated at 60°C for 1 hour. Following which lungs were re-hydrated through an ethanol gradient (95%-70%-50%) and water for 2 hours each. Rehydrated lungs were lysed using 2 mL of genomic lysis buffer (100 mM Tris-HCl pH8.0, 50 mM EDTA, 1% SDS) containing 1 mg/mL proteinase K (Millipore, cat.#70663-4). Briefly, cells were first homogenized (Cole-Palmer, cat.# UX-04727-89) for 1 minutes followed by a 10 sec sonication (2s ON, 2s OFF cycling) to fully disrupt cell membranes and shear DNA (Qsonica, cat.#Q700A). Cells were next incubated in the lysis/proteinase buffer for 48 hours at 60C. Genomic DNA was purified using established phenol-chloroform extraction methods. Briefly, 4 mL of phenol-chloroform (Fisher Scientific cat.# BP1753I-100) was added to the lyzed cells and vortexed at max speed for 2 minutes, followed by spinning tubes at 7000xg for 10 min. The aqueous phase (∼2 mL) containing DNA was carefully extracted, 200 uL of 3 M sodium acetate and 5 mL of 70% ethanol was added to samples and vortexed for 2 min. Samples were placed at -20°C overnight to precipitate genomic DNA. The next day, samples were centrifuged at >7,000g for 1 hour to pellet genomic DNA. Supernatant was carefully aspirated, samples were washed twice with 10 mL of 70% ethanol, vortexed and pelleted at >7,000g for 15 min. Ethanol was aspirated, samples were air-dried for 10-15 minutes until all ethanol had visibly evaporated. Genomic DNA pellet was resuspended in 250-500 uL of pre-warmed (65C) resuspension buffer (5 mM Tris-HCl, pH8.5).

### Western blotting

A-549 cells were treated or not with 5 ng/mL of TGFβ to induced epithelial-mesenchymal transition. After 48 h, cells were lysed in RIPA buffer (0.1% sodium dodecyl sulphate, 50 mM Tris pH 7.4, 150 mM NaCl, 1 mM EDTA, 1% Triton X-100, 1% deoxycholic acid, 200 μM Na_3_VO_4_, and 1mM NaF) supplemented with HALT^TM^ Protease and Phosphatase Inhibitor Cocktail (Thermo Fisher). Lysates were sonicated prior to centrifugation at 4°C for 20 min at 13,000 rpm. Protein concentration was quantified using the DC protein assay (Bio-Rad Laboratories) and a standard curve obtained with increasing concentrations of Bovine Serum Albumin (BSA; Fisher Scientific). Protein samples were normalized to 20 μg total protein per lane and resolved by SDS-PAGE electrophoresis. 4X Bolt LDS sample buffer supplemented with 400 mM DTT (final concentration 1X Bolt LDS with 100 mM DTT) was added to samples before heating at 70°C for 10 min. SDS-PAGE was run using NuPAGE Bis-Tris 4%-12% gradient gels (Thermo Fisher) with MOPS running buffer according to manufacturer’s instructions. Spectra™ Multicolor Protein Ladder (Thermo Fisher) was included as protein ladder on the same gel. Proteins were transferred to a PVDF membrane using the Bio-Rad Turbo Transblot System and transfer kit (Bio-Rad Laboratories) and fixed in methanol prior to blocking in 5% non-fat milk in TNT buffer (50 mM Tris pH 8.0, 150 mM NaCl, 0.1% Tween 20) for 2 h at room temperature. Primary antibody labeling was performed overnight at 4°C using either mouse anti-FLAG M2 epitope tag HRP conjugate (Sigma, cat#A8592) or the following primary antibodies diluted in TNT buffer containing 3% BSA: mouse anti-E-cadherin and anti-N-cadherin (BD Biosciences), rabbit anti-GAPDH (Cell Signaling Technology). Membranes were washed three times prior to secondary antibody labeling for 1 h at room temperature with goat secondary antibodies against mouse or rabbit IgG conjugated to HRP (MilliporeSigma; DC02L and SAB3700878). SuperSignal™ West Pico PLUS Chemiluminescent Substrate (Thermo Fisher) were used to detect HRP-conjugated secondary antibodies according to manufacturer’s instructions prior to imaging. Membranes were imaged on a ChemiDoc™ imaging system and analyzed with Image Lab software (Bio-Rad Laboratories). When necessary, membranes were re-blotted using Re-Blot Plus Strong Solution (MilliporeSigma) for 1 h before blocking and immunoblotting with different primary antibodies.

### Immunofluorescence

A-549 and HaCaT cells were plated at 30-40% confluency in a 96-well format and treated or not with 5 ng/mL of TGFβ and/or 5 μM SB431542, a specific inhibitor of the TGFβ Type I receptor kinase activity, for 48 and 72 h, respectively. DMSO was added as vehicle in the control wells. Cells were washed once with warm PBS before fixing in 4 % paraformaldehyde for 20 min at room temperature, washed twice, and held in PBS at 4°C until immunostaining was conducted. Cells were permeabilized and blocked in 5% normal goat serum (Fisher Scientific) and 0.1% Triton X-100 in PBS for 2 h at room temperature. Primary antibody rabbit anti-Fibronectin (MilliporeSigma) was diluted in blocking buffer and labeling was performed overnight at 4°C. Cells were washed six times with PBS (2 x quick, 2 x 10 min, 2 x 5 min) prior to incubating with goat anti-rabbit secondary antibodies conjugated to Alexa Fluor 555 (Thermo Fisher) for 1 h at room temperature. Cells were then washed six times with PBS as described above, and DAPI (Thermo Fisher) was included in the third wash to counterstain nuclei. Automated full-well imaging was performed using an ImageXpress Pico Cell Imaging System (Molecular Devices) to visualize changes in fibronectin expression between treatment groups. Analysis was performed using the MetaXpress software to determine ranges for fluorescence intensity based on thresholds for minimum and maximum pixel intensities. DAPI staining was included in the analysis to detect and count individual cells.

### Luciferase reporter assay

Basal CREB activity in UM-HMC-1, UM-HMC-3A, and UM-HMC-3B cells was assessed using the Cre-Luc luciferase reporter assay. Briefly, 100,000 cells of each line were seeded per well in a 12-well plate one day prior to transfection. Four hours before transfection, the medium was replaced with antibiotic-free medium. Each well (in duplicate or triplicate) was co-transfected with 1 µg of Cre-Luc luciferase reporter plasmid (p*Cre*-Luc, Clonetech, K2049-1) and 100 ng of the mCherry expression plasmid (pSport6-mCherry, generated in-house) using Lipofectamine 3000 transfection reagent (Thermo Scientific). For the no-reporter control, cells were transfected with 100 ng of pSport6-mCherry together with 1 µg of pcDNA3.1 empty vector. Eighteen hours after transfection, fresh medium was added. Forty-eight hours post-transfection, cell lysates were prepared in 200 µL of BrightGlo Luciferase Assay Buffer (Promega kit-E2650). From each lysate, 50 µL was transferred to an opaque white 96-well plate. mCherry fluorescence was measured first, followed by luciferase activity after adding 50 µL of BrightGlo Luciferase substrate (Promega kit E2650), using the Cytation 5 Hybrid Multi-Mode Reader (BioTek). Luciferase activity was normalized to transfection efficiency based on mCherry fluorescence. The normalized activity for each cell line was expressed as the ratio of reporter-transfected cells to no-reporter controls.

### Flow cytometry

The cellular plasticity associated with EMT and hybrid E/M states was analyzed in parental MEC cell lines (UM-HMC-1, UM-HMC-3A and UM-HMC-3B), UM-HMC-1 cells with A-CREB overexpression, PC-3 cells with VGLL3 knockdown or with KLF3 overexpression, and A-549 cells with KLF3 overexpression, using flow cytometry. EMT markers were selected based on studies by Kröger et al. (PNAS, 2019)^42^, Pastushenko et al. (Nature, 2019)^8^, and are described in Fig. 3g and Extended Fig. 5p. In UM-HMC-1 cells, A-CREB overexpression was induced using doxycycline at concentrations of 250, 500, 750, and 1000 ng/mL for 72 hours. EMT induction in PC-3 cells (VGLL3 knockdown or KLF3 overexpression) was achieved by treating cells with 5 ng/mL or 7.5 ng/mL TGFβ for 72 hours. In A-549 cells with KLF3 overexpression, EMT was induced using 5 ng/mL TGFβ for 72 hours. To counteract TGFβ effects, PC3 and A-549 cells were also treated with 5 μM of the TGFβ inhibitor (SB431542). Following treatment, cells were detached using Accutase Cell Dissociation Solution (Sigma-Aldrich), washed with 1× PBS, and stained with Zombie UV dye for 15 minutes at room temperature in the dark to assess viability. After washing, cells were incubated with 200 µL of extracellular antibody mix in flow buffer (1% BSA/PBS) for 15 minutes at room temperature in the dark. Cells were then washed again using flow buffer, fixed and permeabilized in 125 µL of a 1:5 dilution of Cytofix/Cytoperm solution (BD Biosciences, cat#554722) for 15 minutes at room temperature. Subsequently, 100 µL of a 1:200 dilution of Vimentin antibody was added, and cells were incubated for another 15 minutes at room temperature. After a final wash, cells were resuspended in 500 µL of flow buffer and analyzed using an LSRII flow cytometer (BD Biosciences). Flow cytometry data were processed using FlowJo software (Tree Star). Cell populations were gated based on appropriate unstained and fluorescence-minus-one (FMO) controls.

Extracellular marker antibodies: Alexa Fluor 700 anti-Human CD326/EpCAM (1:400 cat#324243), APC/Fire 750 anti-Human CD24 (1:400, cat#311139), FITC anti-Human CD44 (1:400, cat#397517), and APC anti-Human CD51 (1:400, cat#327913) were purchased from Biolegend. BV510 anti-Human CD61 (1:400, cat#563303), and BD Optibuild BUV737 anti-Human CD104 (1:400, cat#749023) were purchased from BD Biosciences. PE-Vio 615 anti-Human CD106 (1:200, cat#130-129-377) was purchased from Miltenyi Biotech. Intracellular marker antibody: PE anti-Human Vimentin (1:400, BD Biosciences, cat#562337).Live-dead viability dye: Zombie Violet Live-Dead (Biologend, cat#423107).

### Bioinformatics

Bulk RNA-sequencing was performed on either the UM-HMC-1 and UM-HMC- 3A cells expressing inducible A-CREB, or on the PanCancer cell lines (A-549, PC-3, MDA-MB- 231, A-498, and T-24) expressing shRNAs for either control (shNS) or *VGLL3* knockdown (shVGLL3_#2). For the UM-HMC lines, cells were seeded into 6 well plates with medium +/- Dox. Fresh Dox was added every 24 hours. Cells were split 72 hours post seeded and re- seeded into fresh plates. For the PanCancer cell lines, cells were utilized at early passages (<5 passages) following transduction with the respective shRNA viruses. For knockdown + TGFβ stimulated conditions (PC-3), cells were treated with 5 ng/mL TGFβ (Sigma-Aldrich cat.#T7039) 24 hours post-seeding and 72 hours post TGFβ addition cells were isolated for RNA isolation. Cells were trypsinized, pelleted, and lyzed for RNA extraction at the indicated time points and mRNA was extracted as described above. Cells were treated as indicated, cells were lyzed, and RNA was extracted using NucleoSpin RNA kit (Macherey-Nagel, #740955) according to the manufacturer’s protocol. RNA sequencing libraries were prepared at the UNC-TGL core facility using a Bravo Automated Liquid-Handling Platform (Agilent G5562A). The TruSeq Stranded Total RNA Library Prep Ribo-Gold Kit (Illumina 20020599) was used to prepare the RNA libraries so as to deplete contaminating and undesired ribosomal RNA carryover, according to the manufacturer’s protocol (Illumina 1000000040499). Following which RNAseq library quality and quantity were quantified using a TapeStation 4200 (Agilent G2991AA). Quantified and QC’ed libraries were pooled at equal molar ratios and denatured (Illumina 15050107). RNA libraries were sequenced at the UNC-High Throughput Sequencing Facility, wherein all biologic replicates were for the various conditions (12 conditions, in biologic replicates) were sequenced one 1 lane of a NoveSeq6000 S1 flow cell (2 lanes were used for all 24 samples combined). with 2x150 bp paired-end configuration according to the manufacturer’s protocol.

The fastq files were aligned to the GRCh38 human genome (GRCh38.d1.vd1.fa) using STAR v2.4.2^43^ with the following parameters: --outSAMtype BAM Unsorted --quantMode TranscriptomeSAM. Transcript abundance for each sample was estimated with salmon v0.1.19^44^ to quantify the transcriptome defined by Gencode v22. Gene level counts were summed across isoforms and genes with low counts (maximum expression < 10) were filtered for the downstream analyses. We tested genes for differential expression (p.adj < 0.01, fold change > 1.5 and baseMean > 10) in DESeq2 v1.38.2^45^ in R and gene set variation analysis on Molecular Signatures Database (MsigDB)^46^ C2 curated gene sets^47^ was conducted with GSVA v1.30.1^48^ in R where cell line is included as a covariate.

Batch effects between cell lines were removed using removeBatchEffect function in limma v3.46.0. Unsupervised and supervised hierarchical clustering, dendrogram generation, and heat-map plotting were performed with ComplexHeatmap v2.6.2^49,50^. A complete linkage algorithm with Euclidean distance function was applied to batch-corrected, vst normalized expression data for statistically significant differentially expressed genes. Principal component analysis was performed using the ‘prcomp’ function in R, in which a singular value decomposition algorithm was applied to a centered and scaled correlation matrix of batch-corrected and vst normalized gene expression data.

DEG analyses were performed comparing A-CREB-Day5 to Non-treated samples in UM- HMC-1 and UM-HMC-3A cell lines independently. Genes with p.adj < 0.01, fold change > 1.5 and baseMean > 10 were called as differentially expressed genes (DEGs) and the overlap of DEGs between two cell lines were defined as A-CREB gene signature. A subset of genes in A-CREB signature was selected to construct A-CREB_Full EMT signature if a gene meets one of the following criteria: 1) overlap of DEGs in at least 4 out of 12 EMT experiments with same directionality (i.e. both up-regulated upon A-CREB treatment and induced EMT treatment)^16^, 2) overlap with same MsigDB Hallmark EMT gene set; 3) overlap with same directionality to PanCancer EMT^5^. The A-CREB_NoEMT signature was derived in a similar fashion such that genes overlapping with any of the following gene sets were removed from the A-CREB signature: 1) overlap of DEGs in at least 4 out of 12 EMT experiments with same directionality (i.e. both up-regulated upon A-CREB treatment and induced EMT treatment)^16^, 2) overlap with same MsigDB Hallmark EMT gene set; 3) overlap with same directionality to PanCancer EMT^5^.

Samples were categorized into four groups based on mean expression of genes in A-CREB_Full EMT, A-CREB_EMT, or A-CREB_NoEMT gene signatures. Kaplan-Meier plots were generated to compare progression free survival (PFS) patterns between the quartiles with highest and lowest expression level of signature of interest. Differences in PFS patterns were examined using log-rank tests. Prognostic value of selective genes were evaluated using univariate Cox proportional hazards model to compute hazard ratios. All survival analyses were performed with R packages survival v3.2.7, survminer v0.4.9 and forestmodel v0.6.2.

For *VGLL3* and *KLF3* promoter analyses, all putative transcription factor binding sites as defined by the JASPAR CORE^51^ with a p-value <= 10^-5^ were reported for the 2.5kb upstream and 500bp downstream of the human *VGLL3* and *KLF3* transcription start sites.

For scRNA-seq pseudotime analysis, counts tables and metadata from time course experiments in Cook et. al. were downloaded from GEO (GSE147405)^16^. All cells provided on GEO were utilized in statistical analyses. A generalized additive model (∼ s(pseudotime) + batch) was utilized via the mgcv package (v1.8-42)^52^ to test for an association between pseudotime and *VGLL3* expression as in Cook et al., with k=4 basis functions; smoothing parameters were selected via restricted maximum likelihood (REML). Log normalized counts were used. P-values for GAM models were adjusted for multiple comparisons using the Bonferroni correction; tests were not conducted for genes with fewer than 10 counts total. Additionally, cell level counts were aggregated to pseudobulk counts at the sample level and a likelihood ratio test was conducted in DESeq2 (v1.38.0)^45^ in R (v4.2.2) to test for an association between treatment time and expression. Genes with fewer than 10 counts in at least one sample were excluded from analyses. P-values for likelihood ratio tests were adjusted for multiple comparisons utilizing the Benjamini-Hoechberg procedure.

---

**Extended Data Figure S1. a-b,** Interactive cluster-grams for ChEA3 enrichment analyses of CREB/ATF1 family transcription factors and differentially expressed genes related to canonical (**a**) MSigDB Hallmark EMT or (**b**) Tan et al. Pan-Cancer derived EMT gene lists. **c-e,** Pie charts depicting the percentage of genes within each corresponding EMT gene signature predicted to be associated with regulation by one or more CREB/ATF1 family of transcription factors.

**Extended Data Figure S2. Generation and validation of CREB-dependent cell line models expressing an inducible dominant negative CREB molecule, A-CREB. a,** Diagram of the recurrent balanced chromosomal translocation present in ≥50% of salivary mucoepidermoid carcinomas and the transcriptional coactivators CRTC1 and MAML2 involved in the resulting gene fusion. **b,** Diagram highlighting the mechanism whereby A-CREB disrupts gene expression. *Top*, CRTC1-MAML2 (C1/M2) regulates adaptive response gene expression via direct binding between the CREB binding domain of CRTC1 to the bZip domain of the transcription factor CREB. *Bottom*, A-CREB dimerizes with endogenous CREB and the acidic domain of A-CREB prevents binding of CREB to cAMP response element (CRE) motifs within the promoters of adaptive response genes thus attenuating gene expression. **c,** Representative western blot validation of Dox-inducible FLAG-epitope tagged A-CREB expression (*n* = 3). **d,** Representative immunofluorescence validation of FLAG-epitope tagged A-CREB expression and nuclear localization following Dox addition (*n* = 2). **e,** qPCR validation of sustained CREB- target gene inhibition following Dox-induced expression of A-CREB (*n* = 3, mean ± SEM, **** *p* < 0.0001).

**Extended Data Figure S3. Phenotypic characterization of tumor cell behavior after CREB disruption. a-b,** Quantification of relative cell proliferation (**a**) and relative cell viability (**b**) observed post-Dox addition to induce A-CREB versus control non-treated UM-HMC-1 and UM- HMC-3A cell lines (*n* = 3, mean ± SEM, **** *p* < 0.0001). **c,** Representative transwell invasion assays for UM-HMC-1 and UM-HMC-3A cell lines with or without induced A-CREB expression 72 hours prior. **d,** Representative image of the surgical procedure optimized for orthotopic salivary gland xenografts. The black arrow outside the green circle annotation highlights the location of an incision made to expose the murine submandibular gland into which the UM-HMC cell lines mixed with Matrigel were injected. **e,** Schematic of the orthotopic salivary gland (SMG) xenograft studies designed to assess the effects of disrupting CREB activity on E/M plasticity and *in vivo* metastatic potential. Animals were maintained on a normal chow and then group A remained on normal chow while groups B and C were switched to a Dox chow once the average tumor size reached 125 mm^3^. The Dox chow was maintained until Day 14 for group B and until Day 21 for group C at which time the Dox chow was withdrawn and switched back to normal chow. Animals were sacrificed when tumors reached ∼1,000 mm^3^.

**Extended Data Figure S4. CREB binding motif enrichment and gating strategy for flow cytometry analysis of EpCam^+^ and Vimentin^+^ cell populations. a,** CREB-related transcription factor motif enrichment in ATAC-seq datasets representative of EMT transition states as determined by Homer analysis by Pastushenko et al. Enrichment of the CREB5 motif was determined for peaks that were upregulated between the indicated subpopulations in cells undergoing (**a**) EMT or (**b**) MET. **c,** Flow plots outlining the gating strategy employed to identify viable EpCam+/- and Vimentin +/- cells for further hybrid E/M state subpopulation analysis. **d,** Summary marginal flow plots with expression profiles showing the basal EpCam and Vimentin expression levels according to CREB activity levels for the UM-HMC-1 (CREB^hi^), UM-HMC-3A (CREB^mid^), and UM-HMC-3B (CREB^lo^) cell lines combined (*left* plot) or as each individual cell line.

**Extended Data Figure S5. Gating strategy for flow analysis of the different hybrid E/M cell populations in the EpCam^+/-^ and Vimentin^+/-^ subsets. a,** Proportion of cells expressing EpCam (Ep+/-) and/or Vimentin (Vim+/-) for each UM-HMC cell line. **b-d,** Flow cytometry profiles showing the six different hybrid E/M populations based on expression of CD106, CD51, and CD61 expression in the (**b**) UM-HMC-1 (CREB^hi^), (**c**) UM-HMC-3A (CREB^mid^), and (**d**) UM- HMC-3B (CREB^lo^) cell lines. **e-g,** Percentage of EpCam^+/-^ cells within each hybrid E/M population for (**e**) all viable EpCam^+^ and EpCam^-^ cells, (**f**) only viable EpCam^+^ cells, or (**g**) only viable EpCam^-^ cells. (**h-o**) Flow plots outlining the gating strategy employed to establish early versus late hybrid E/M states based on the expression of CD24, CD51, and CDE104 in viable EpCam^+^ and EpCam^-^ cells. (**p**) Diagram depicting the expression for a condensed subset of cell surface markers present on epithelial, hybrid E/M, and mesenchymal tumor cells. Early hybrid E/M cell populations are CD24^lo^/CD104^lo^/CD51^lo^ and overlap with TN and SP1 hybrid E/M cells, while late hybrid E/M cells are CD24^lo^/CD104^hi^/CD51^hi^ and overlap with SP2, DP1, DP2, and TP hybrid E/M cells.

**Extended Data Figure S6. PanCancer effects of regulating CREB activity. a,** Quantification of relative cell proliferation observed post-Dox addition to induce A-CREB versus control non- treated UM-HMC-1 and UM-HMC-3A cell lines (*n* = 3, mean ± SEM, \**p* < 0.05, \*\*\*\**p* < 0.0001). **b,** Time-lapse quantification of collective cell migration/scratch closure for various cell lines with or without A-CREB expression 72 hours prior (*n* = 3, mean ± SEM plotted within error bars as shaded region, \**p* < 0.05, \*\*\**p* < 0.001). **c,** Kaplan Meier patient survival plots for the four TCGA cancers that display significant HR according to the top and bottom quartiles of expression for each respective signature. Significance was determined using a log-rank test (\**p* < 0.05, \*\**p* < 0.01, \*\*\**p* < 0.001, \*\*\*\**p* < 0.0001).

**Extended Data Figure S7. PanCancer association of non-canonical A-CREB-NoEMT genes with Hallmark_EMT and regulators of epithelial versus mesenchymal cell state*s*. a,** PanCancer correlation of gene expression for shared non-canonical A-CREB_NoEMT core genes *KLK7*, *KRT16*, *IL1RN*, *MARCO*, *PTGS2,* and *VGLL3* with the Hallmark_EMT gene signature. Pearson correlation plotted with respect to significance where each dot represents a unique cancer type across 33 TCGA cancers tested. Black dots are not significant and pink dots are significant (p < 0.05). **b,** Scatter plots showing the correlation of *VGLL3* gene expression with gene expression for several key canonical regulators of epithelial versus mesenchymal cell states. Spearman correlation was performed between *VGLL3* and the TCGA Breast Invasive Carcinoma PanCancer Atlas (*n* = 944) with *p*-value and R^2^ values displayed indicating a negative correlation with epithelial state drivers and a positive correlation of *VGLL3* expression with mesenchymal state drivers.

**Extended Data Figure S8. PanCancer validation of VGLL3 as a *bona fide* orchestrator of EMT. a-l,** Evaluation of *VGLL3* knockdown on six different cell lines with high basal EMT- associated invasiveness. Two independent shRNAs targeting *VGLL3* were examined in stable knockdown cells representative of (**a**) pancreatic, (**c, g**) renal, bladder, (**e**) bladder, (**i**) head and neck, and (**k**) ovarian cancers for their effects on gene expression by real-time qPCR of endogenous *VGLL3* expression levels. Fold expression is shown normalized to *RPL23* and relative to the no Dox control cells (*n* = 3, mean ± SEM, * *p* < 0.05, *** *p* < 0.001, **** *p* < 0.0001). Representative images and quantification of transwell assays in stable *VGLL3* knockdown cells representative of (**b**) pancreatic, (**d, h**) renal, bladder, (**f**) bladder, (**j**) head and neck, and (**l**) ovarian cancers for effects of knockdown on invasion. Chambers with 8 μm pores were coated with 5% Matrigel prior to cell seeding (*n* = 3, mean ± SEM, * *p* < 0.05, ** *p* < 0.01, *** *p* < 0.001, **** *p* < 0.0001).

**Extended Data Figure S9. *VGLL3* controls metastatic behavior *in vitro* and *in vivo*. a,** Representative images and quantification of transwell assays in A-549 cells with stable *VGLL3* knockdown examined for effects on invasion. Chambers with 8 μm pores were coated with 5% Matrigel prior to cell seeding (*n* = 3, mean ± SEM, **** *p* < 0.0001). **b,** Time-lapse quantification of metastatic burden in the lungs of mice receiving tail-vein injections of A-549 cells stably expressing a LumiFluor bioluminescent reporter (GpNLuc) and either control non-specific (shNS) or *VGLL3*-targeting (shVGLL3_#2) shRNAs. Radiance BLI measurements were collected from regions of interest on the time points indicated and plotted (n = 5 per group, mean ± SEM, * *p* < 0.05). **c-d,** Experimental induction of EMT with TGFβ induces *VGLL3* gene expression and is reversible following withdrawal or knockdown. **c,** Sina plot of data collected by Cook et al. showing the distribution of *VGLL3* gene expression across time points for either control A-549 cells or cells treated with TGFβ gene for the indicated length of time prior to withdrawal of TGFβ stimulation. **d,** Heatmap of DEGs between untreated parental PC-3 cells or stable PC-3 cells expressing either control non-specific (shNS) or *VGLL3*-targeting (shVGLL3_#2) shRNAs treated with TGFβ stimulation. Unsupervised hierarchical clustering performed on 653 DEGs with fold change > 1.5, padj < 0.01, and base mean > 10 (*n* = 2 per treatment/group). **e,** ACREB_Full EMT signature enrichment scores in the indicated stable cell lines expressing either control non-specific (shNS) or *VGLL3*-targeting (shVGLL3_#2) shRNAs (*n* = 2 per group).

**Extended Data Figure S10. Identification and characterization of the transcriptional repressor *KLF3* as a CREB regulated target controlling epithelial states. a,** HaCaT cells were treated with or without 5 ng/mL of TGFβ and/or 5 μM SB431542 for 72 h before fixation and immunofluorescence staining using an antibody against fibronectin. Cells were analyzed using an ImageXpress Pico Cell Imaging System, and the average fluorescence intensity in each well was determined using MetaXpress software (n = 6, **** *p* ≤ 0.0001, ns = not significant). **b,** Gating strategy for flow cytometry analysis of the different hybrid E/M cell populations in the EpCam^+/-^ and Vimentin^+/-^ subsets for PC-3 cells. The flow cytometry profile shows the six different hybrid E/M populations based on expression of CD106, CD51, and CD61 expression. **c,** Heatmap of DEGs between the indicated cell lines for untreated parental or stable cells expressing either control non-specific (shNS) or *VGLL3*-targeting (shVGLL3_#2) shRNAs and/or treated with TGFβ stimulation. Unsupervised hierarchical clustering of overlapping DEGs performed with genes listed as up or down if they have significantly increased/decreased expression (fold change > 1.5, padj < 0.01) in the PC-3 shNS versus shVGLL3_#2 knockdown and up in all other cell lines indicated (*n* = 2 per treatment/group). **d,** Visualization of UCSC Genome Browser search results on hg38 (Human Dec. 2013 (GRCh38/hg38)) for *KLF3* tracks related to H3K27Ac histone marks of active transcription as determined by a ChIP-seq assay, transcription factor binding (TF rPeak clusters) displayed as clusters of representative transcription factor binding sites identified by ChIP-seq experiments with locations of a cluster of CREB1 sites located downstream of the TSS highlighted, and also multiple alignments of 100 vertebrate species highlighting evolutionary conservation of the region overlapping the cluster of CREB1 binding. **e,** Real-time qPCR of parental A-549 cells and A-549 cells engineered to stably overexpress un-tagged or FLAG epitope-tagged *KLF3*. Fold expression is shown normalized to *RPL23* and relative to the parental cells (*n* = 3, mean ± SEM, **** *p* < 0.0001). **f,** Western blot validation of KLF3 over expression from whole cell lysates probed with either anti-KLF3 or anti-Actin. **g,** Time-lapse cell proliferation assays comparing the parental and *KLF3* stably overexpressing A-549 cell lines. **h,** Quantification of transwell assays in A-549 cells with stable *KLF3* overexpression examined for effects on invasion. Chambers with 8 μm pores were coated with 5% Matrigel prior to cell seeding. Fold change in invasion is shown relative to the parental cells (*n* = 3, mean ± SEM, * *p* < 0.05, ** *p* < 0.01). **i,** Parental and *KLF3* stably overexpressing A-549 cell lines were treated with 5 ng/mL of TGFβ for 48 h, and cell lysates were analyzed by immunoblotting for E-cadherin, N-cadherin, and GAPDH as loading control. **j-k,** Quantification of western blot results presented in panel j for either (**j**) E- cadherin or (**k**) N-cadherin (n = 3, mean ± SD, *** *p*≤0.001, **** *p*≤0.0001, ns: not significant). **l,** Gating strategy for flow cytometry analysis of the different hybrid E/M cell populations in the EpCam^+/-^ and Vimentin^+/-^ subsets for A-549 cells. The flow cytometry profile shows the six different hybrid E/M populations based on expression of CD106, CD51, and CD61 expression.

## References

1 Kalluri, R. & Weinberg, R. A. The basics of epithelial-mesenchymal transition. J Clin Invest 119, 1420–1428 (2009). 10.1172/JCI39104

2 Fleischmajer, R., Billingham, R. E., Hahnemann Medical, C. & Hospital of, P. Epithelial-mesenchymal interactions; 18th Hahnemann symposium. (Williams & Wilkins, 1968).

3 Hay, E. D. An overview of epithelio-mesenchymal transformation. Acta Anat (Basel) 154, 8–20 (1995). 10.1159/000147748

4 Nieto, M. A., Huang, R. Y., Jackson, R. A. & Thiery, J. P. Emt: 2016. Cell 166, 21–45 (2016). 10.1016/j.cell.2016.06.028

5 Tan, T. Z. et al. Epithelial-mesenchymal transition spectrum quantification and its efficacy in deciphering survival and drug responses of cancer patients. EMBO Mol Med 6, 1279–1293 (2014). 10.15252/emmm.201404208

6 Yang, J. et al. Guidelines and definitions for research on epithelial-mesenchymal transition. Nat Rev Mol Cell Biol 21, 341–352 (2020). 10.1038/s41580-020-0237-9

7 Petersen, O. W. et al. The plasticity of human breast carcinoma cells is more than epithelial to mesenchymal conversion. Breast Cancer Res 3, 213–217 (2001). 10.1186/bcr298

8 Pastushenko, I. et al. Identification of the tumor transition states occurring during EMT. Nature 556, 463–468 (2018). 10.1038/s41586-018-0040-3

9. Puram, S. V., et al. in *Cell* (2017).

10 Pal, A., Barrett, T. F., Paolini, R., Parikh, A. & Puram, S. V. Partial EMT in head and neck cancer biology: a spectrum instead of a switch. Oncogene 40, 5049–5065 (2021). 10.1038/s41388-021-01868-5

11 Nestler, E. J. Cellular responses to chronic treatment with drugs of abuse. Crit Rev Neurobiol 7, 23–39 (1993).

12 Ghosh, A., Ginty, D. D., Bading, H. & Greenberg, M. E. Calcium regulation of gene expression in neuronal cells. J Neurobiol 25, 294–303 (1994). 10.1002/neu.480250309

13 Tasoulas, J., Rodón, L., Kaye, F. J., Montminy, M. & Amelio, A. L. Adaptive Transcriptional Responses by CRTC Coactivators in Cancer. Trends in cancer 5, 111–127 (2019). 10.1016/j.trecan.2018.12.002

14 Srinivasan, S. et al. CREB Drives Acinar to Ductal Cells Reprogramming and Promotes Pancreatic Cancer Progression in Preclinical Models of Alcoholic Pancreatitis. Cell Mol Gastroenterol Hepatol, 101606 (2025). 10.1016/j.jcmgh.2025.101606

15 Mayr, B. & Montminy, M. Transcriptional regulation by the phosphorylation-dependent factor CREB. Nat Rev Mol Cell Biol 2, 599–609 (2001).

16 Cook, D. P. & Vanderhyden, B. C. Context specificity of the EMT transcriptional response. Nature communications 11, 1–9 (2020). 10.1038/s41467-020-16066-2

17 Ahn, S. et al. A dominant-negative inhibitor of CREB reveals that it is a general mediator of stimulus-dependent transcription of c-fos. Molecular and cellular biology 18, 967–977 (1998).

18 Rozenberg, J. M., Bhattacharya, P., Chatterjee, R., Glass, K. & Vinson, C. Combinatorial recruitment of CREB, C/EBPβ and c-Jun determines activation of promoters upon keratinocyte differentiation. PLOS ONE 8, e78179 (2013). 10.1371/journal.pone.0078179

19 Rozenberg, J. et al. Inhibition of CREB function in mouse epidermis reduces papilloma formation. Molecular cancer research : MCR 7, 654–664 (2009). 10.1158/1541-7786.MCR-08-0011

20 Zhang, L. et al. The transcription factor CREB regulates epithelial-mesenchymal transition of lens epithelial cells by phosphorylation-dependent and phosphorylation-independent mechanisms. J Biol Chem 301, 108064 (2025). 10.1016/j.jbc.2024.108064

21 Lamouille, S., Xu, J. & Derynck, R. Molecular mechanisms of epithelial-mesenchymal transition. Nat Rev Mol Cell Biol 15, 178–196 (2014). 10.1038/nrm3758

22 Xie, S. et al. Dominant-negative CREB inhibits tumor growth and metastasis of human melanoma cells. Oncogene (1997).

23 Maiuri, P. et al. The first World Cell Race. Curr Biol 22, R673–675 (2012). 10.1016/j.cub.2012.07.052

24 Mimica, X. et al. Distant metastasis of salivary gland cancer: Incidence, management, and outcomes. Cancer 126, 2153–2162 (2020). 10.1002/cncr.32792

25 Aiello, N. M. et al. EMT Subtype Influences Epithelial Plasticity and Mode of Cell Migration. Dev Cell 45, 681–695 e684 (2018). 10.1016/j.devcel.2018.05.027

26 Aceto, N. et al. Circulating tumor cell clusters are oligoclonal precursors of breast cancer metastasis. Cell 158, 1110–1122 (2014). 10.1016/j.cell.2014.07.013

27 Cheung, K. J. & Ewald, A. J. A collective route to metastasis: Seeding by tumor cell clusters. Science 352, 167–169 (2016). 10.1126/science.aaf6546

28 Hori, N. et al. Vestigial-like family member 3 (VGLL3), a cofactor for TEAD transcription factors, promotes cancer cell proliferation by activating the Hippo pathway. J Biol Chem 295, 8798–8807 (2020). 10.1074/jbc.RA120.012781

29 Yamaguchi, N. Multiple Roles of Vestigial-Like Family Members in Tumor Development. Front Oncol 10, 1266 (2020). 10.3389/fonc.2020.01266

30 Hori, N. et al. Vestigial-like family member 3 stimulates cell motility by inducing high-mobility group AT-hook 2 expression in cancer cells. J Cell Mol Med 26, 2686–2697 (2022). 10.1111/jcmm.17279

31 Xu, J., Lamouille, S. & Derynck, R. TGF-beta-induced epithelial to mesenchymal transition. Cell Res 19, 156–172 (2009). 10.1038/cr.2009.5

32 Katsuno, Y., Lamouille, S. & Derynck, R. TGF-beta signaling and epithelial-mesenchymal transition in cancer progression. Curr Opin Oncol 25, 76–84 (2013). 10.1097/CCO.0b013e32835b6371

33 Jones, J. et al. KLF3 Mediates Epidermal Differentiation through the Epigenomic Writer CBP. iScience 23, 101320 (2020). 10.1016/j.isci.2020.101320

34 Eaton, S. A. et al. A network of Kruppel-like Factors (Klfs). Klf8 is repressed by Klf3 and activated by Klf1 in vivo. J Biol Chem 283, 26937-26947 (2008). 10.1074/jbc.M804831200

35 Funnell, A. P. et al. The CACCC-binding protein KLF3/BKLF represses a subset of KLF1/EKLF target genes and is required for proper erythroid maturation in vivo. Mol Cell Biol 32, 3281–3292 (2012). 10.1128/MCB.00173-12

36 Funnell, A. P. et al. Generation of mice deficient in both KLF3/BKLF and KLF8 reveals a genetic interaction and a role for these factors in embryonic globin gene silencing. Mol Cell Biol 33, 2976–2987 (2013). 10.1128/MCB.00074-13

37 Knights, A. J. et al. Kruppel-like Factor 3 (KLF3/BKLF) Is Required for Widespread Repression of the Inflammatory Modulator Galectin-3 (Lgals3). J Biol Chem 291, 16048–16058 (2016). 10.1074/jbc.M116.715748

38 Warner, K. A. et al. in *Oral Oncol* Vol. 49 1059-1066 (2013).

39 Young, L., Sung, J., Stacey, G. & Masters, J. R. Detection of Mycoplasma in cell cultures. Nature protocols 5, 929–934 (2010). 10.1038/nprot.2010.43

40 Musicant, A. M. et al. CRTC1/MAML2 directs a PGC-1alpha-IGF-1 circuit that confers vulnerability to PPARgamma inhibition. Cell Rep 34, 108768 (2021). 10.1016/j.celrep.2021.108768

41 Amelio, A. L. et al. CRTC1/MAML2 gain-of-function interactions with MYC create a gene signature predictive of cancers with CREB-MYC involvement. Proceedings of the National Academy of Sciences of the United States of America 111, E3260–3268 (2014). 10.1073/pnas.1319176111

42 Kroger, C. et al. Acquisition of a hybrid E/M state is essential for tumorigenicity of basal breast cancer cells. Proc Natl Acad Sci U S A 116, 7353–7362 (2019). 10.1073/pnas.1812876116

43 Dobin, A. et al. in *Bioinformatics* Vol. 29 15–21 (2013).

44 Patro, R., Duggal, G., Love, M. I., Irizarry, R. A. & Kingsford, C. Salmon provides fast and bias-aware quantification of transcript expression. Nature methods 14, 417–419 (2017). 10.1038/nmeth.4197

45 Love, M. I., Huber, W. & Anders, S. Moderated estimation of fold change and dispersion for RNA-seq data with DESeq2. Genome biology 15, 550 (2014). 10.1186/s13059-014-0550-8

46 Mootha, V. K. et al. PGC-1alpha-responsive genes involved in oxidative phosphorylation are coordinately downregulated in human diabetes. Nature genetics 34, 267–273 (2003). 10.1038/ng1180

47 Subramanian, A. et al. in *Proc Natl Acad Sci USA* Vol. 102 15545-15550 (National Academy of Sciences, 2005).

48 Hanzelmann, S., Castelo, R. & Guinney, J. GSVA: gene set variation analysis for microarray and RNA-seq data. BMC Bioinformatics 14, 7 (2013). 10.1186/1471-2105-14-7

49 Chen, Y. et al. A versatile polypharmacology platform promotes cytoprotection and viability of human pluripotent and differentiated cells. Nat Methods 18, 528–541 (2021). 10.1038/s41592-021-01126-2

50 Ritchie, M. E. et al. limma powers differential expression analyses for RNA-sequencing and microarray studies. Nucleic Acids Res 43, e47 (2015). 10.1093/nar/gkv007

51 Castro-Mondragon, J. A. et al. JASPAR 2022: the 9th release of the open-access database of transcription factor binding profiles. Nucleic Acids Res 50, D165–D173 (2022). 10.1093/nar/gkab1113

52 Wood, S. N. Fast Stable Restricted Maximum Likelihood and Marginal Likelihood Estimation of Semiparametric Generalized Linear Models. Journal of the Royal Statistical Society: Statistical Methodology Series B 73, 3–36 (2011). 10.1111/j.1467-9868.2010.00749.x

