## Supplementary figures and images for "Adaptive Responses Directed by CREB Control Epithelial-Mesenchymal Plasticity in Cancer"

### Extended Figure 1

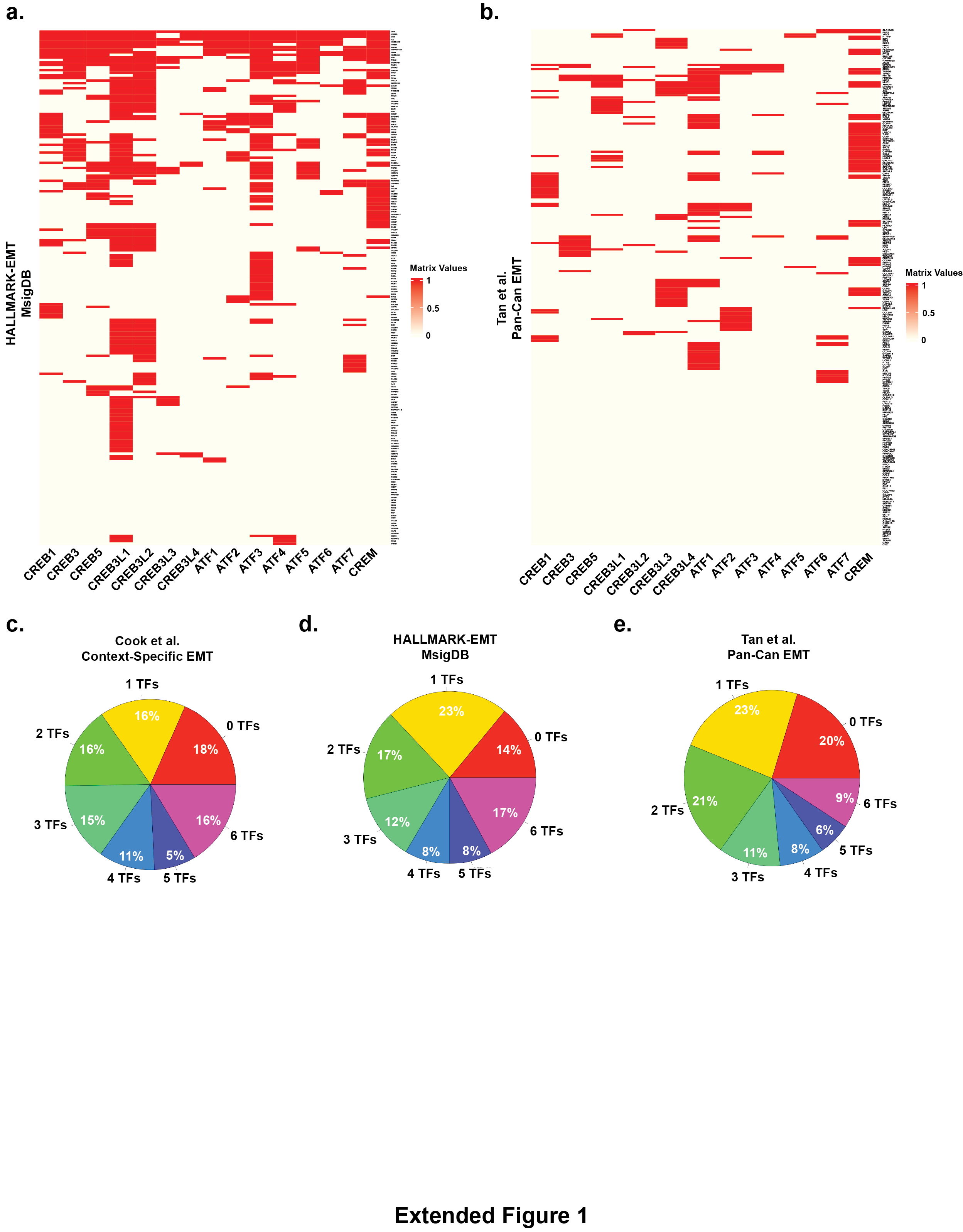

### Extended Figure 2

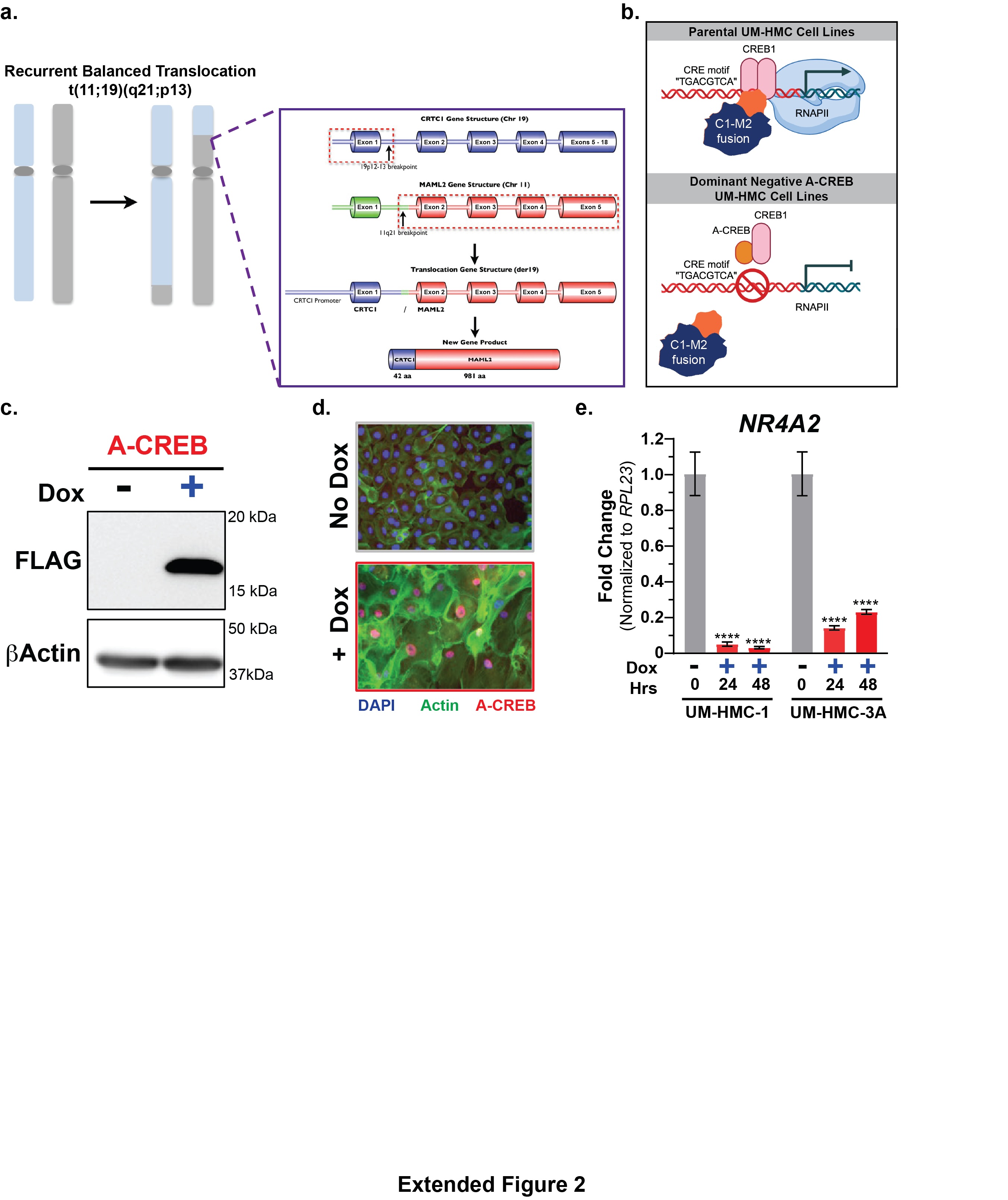

### Extended Figure 3

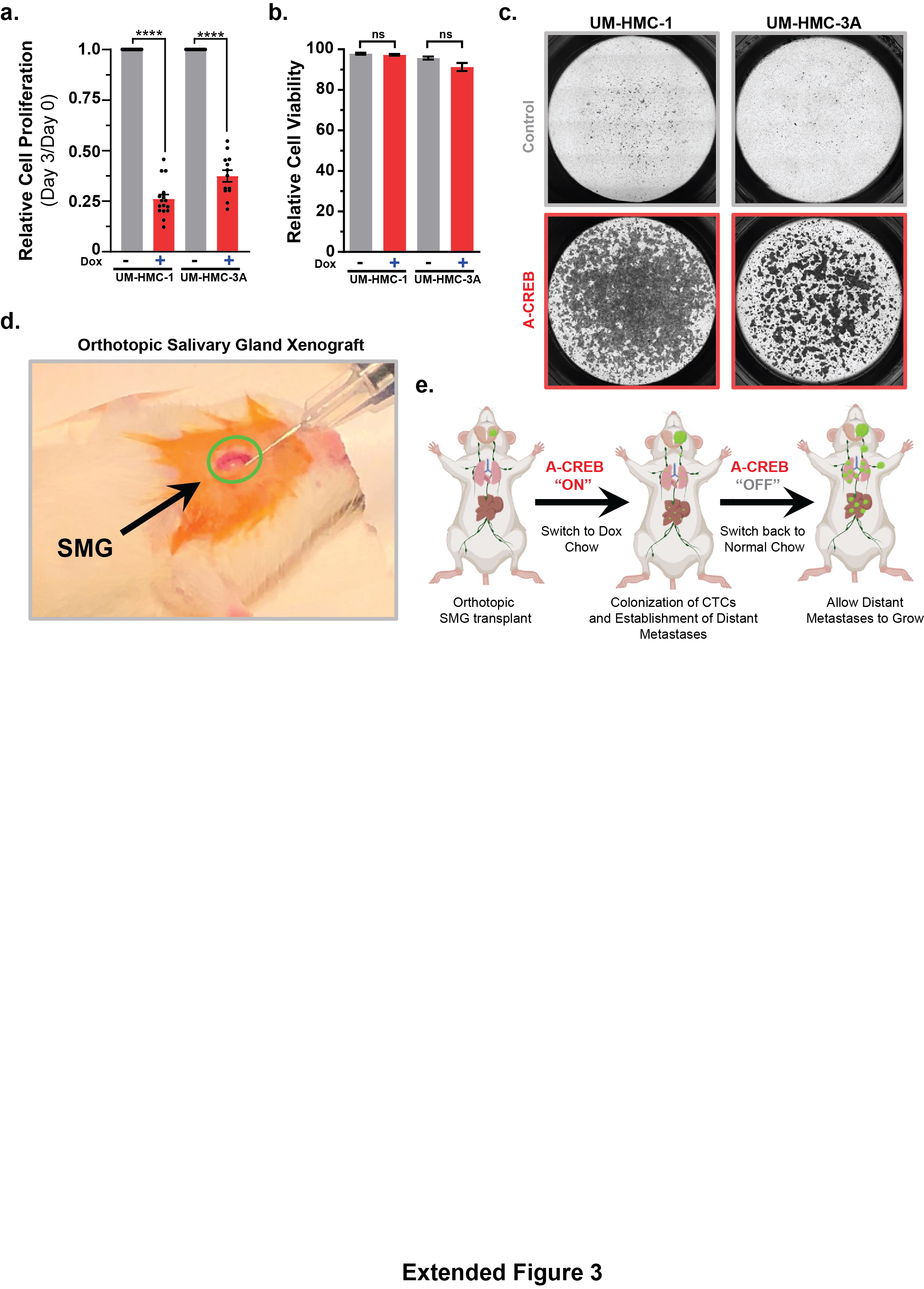

### Extended Figure 4

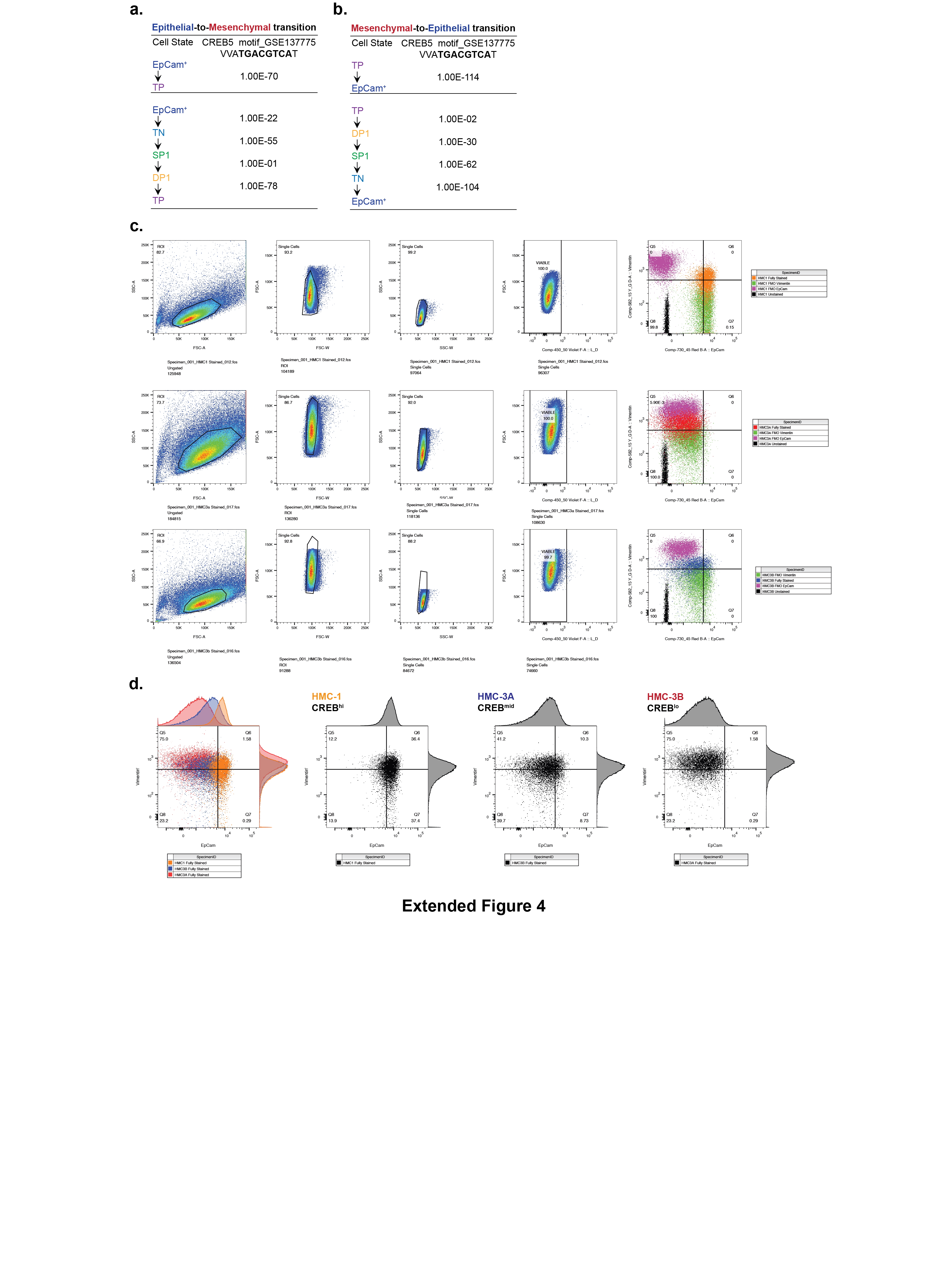

### Extended Figure 5

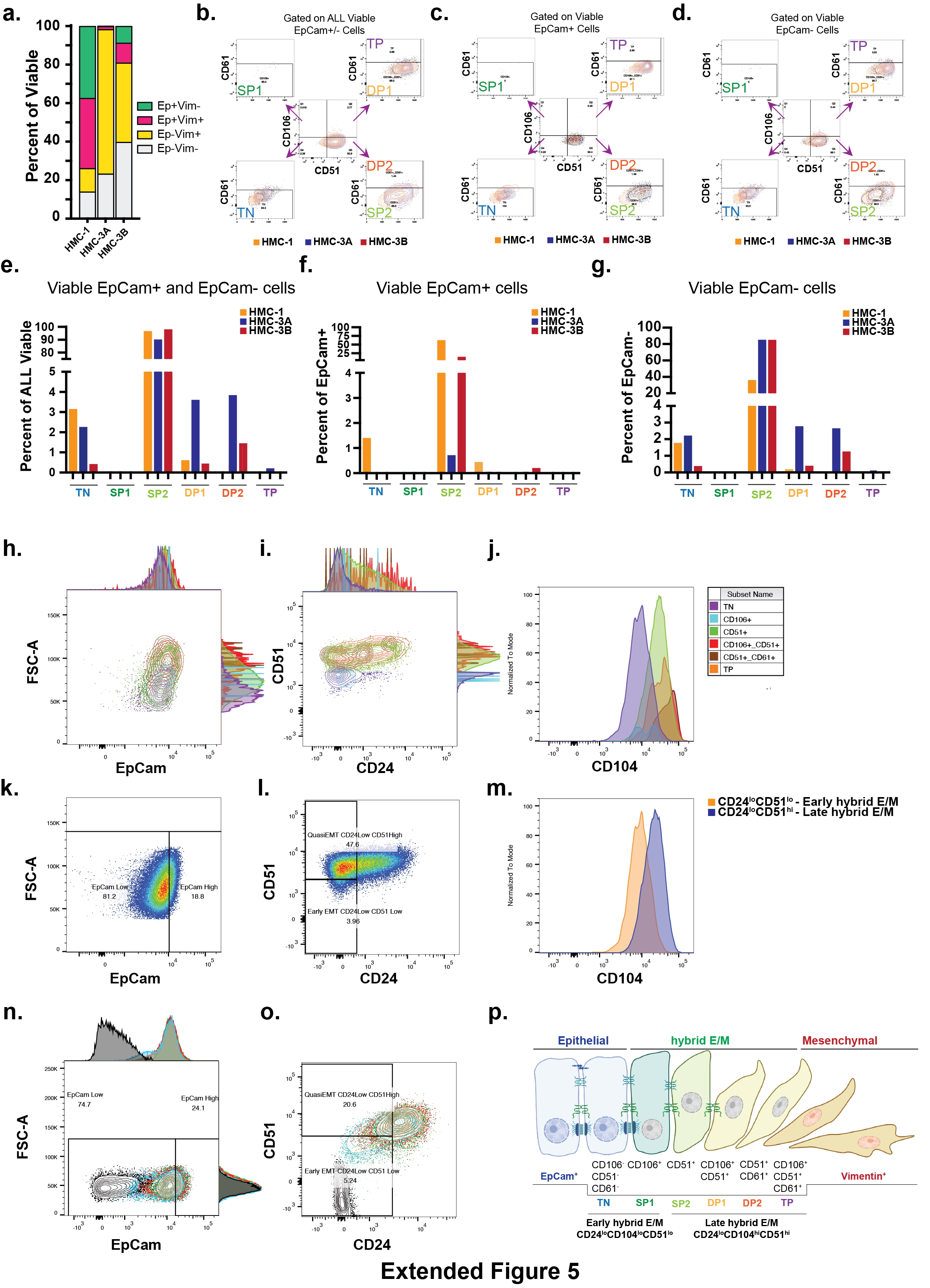

### Extended Figure 6

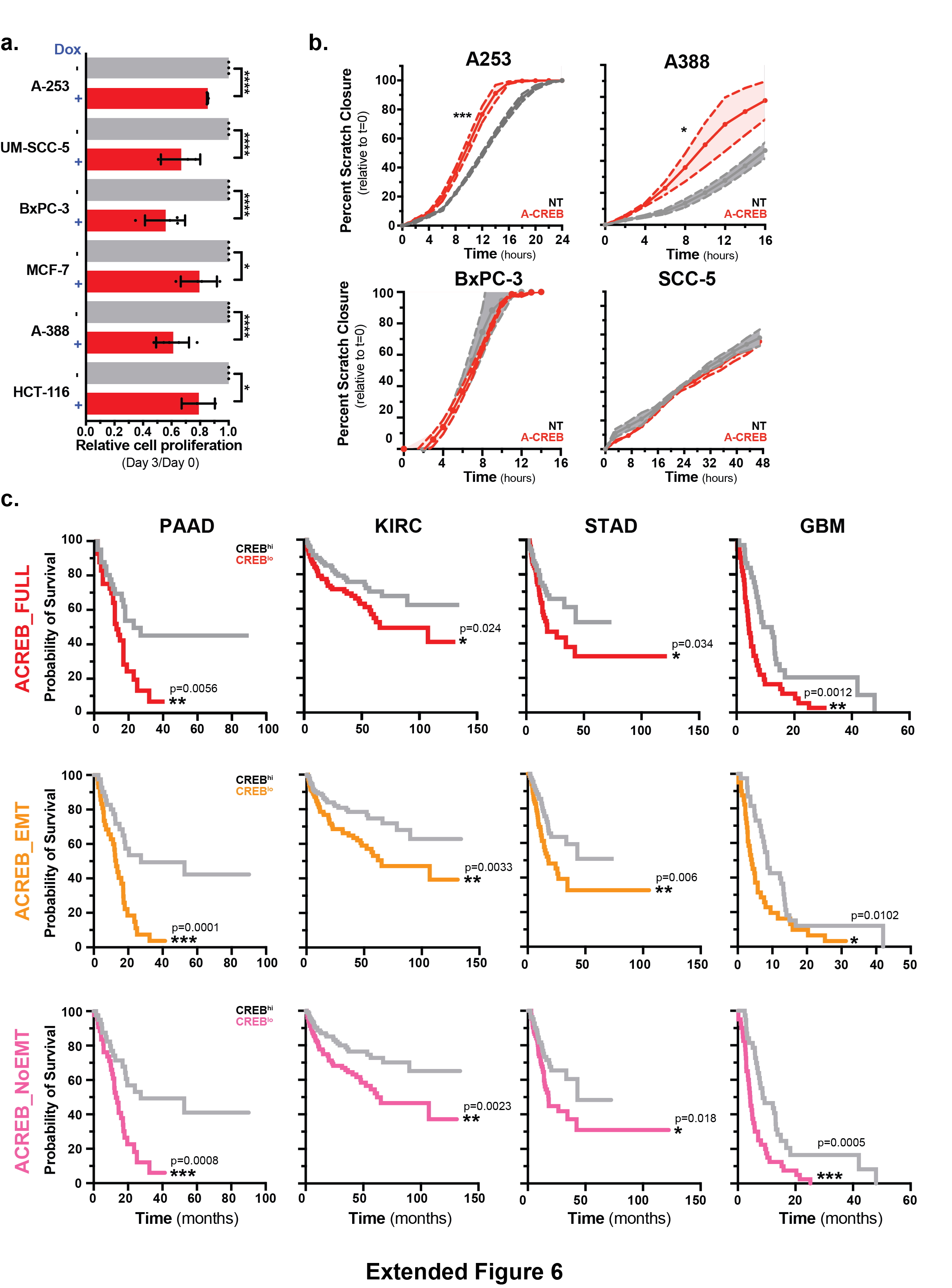

### Extended Figure 7

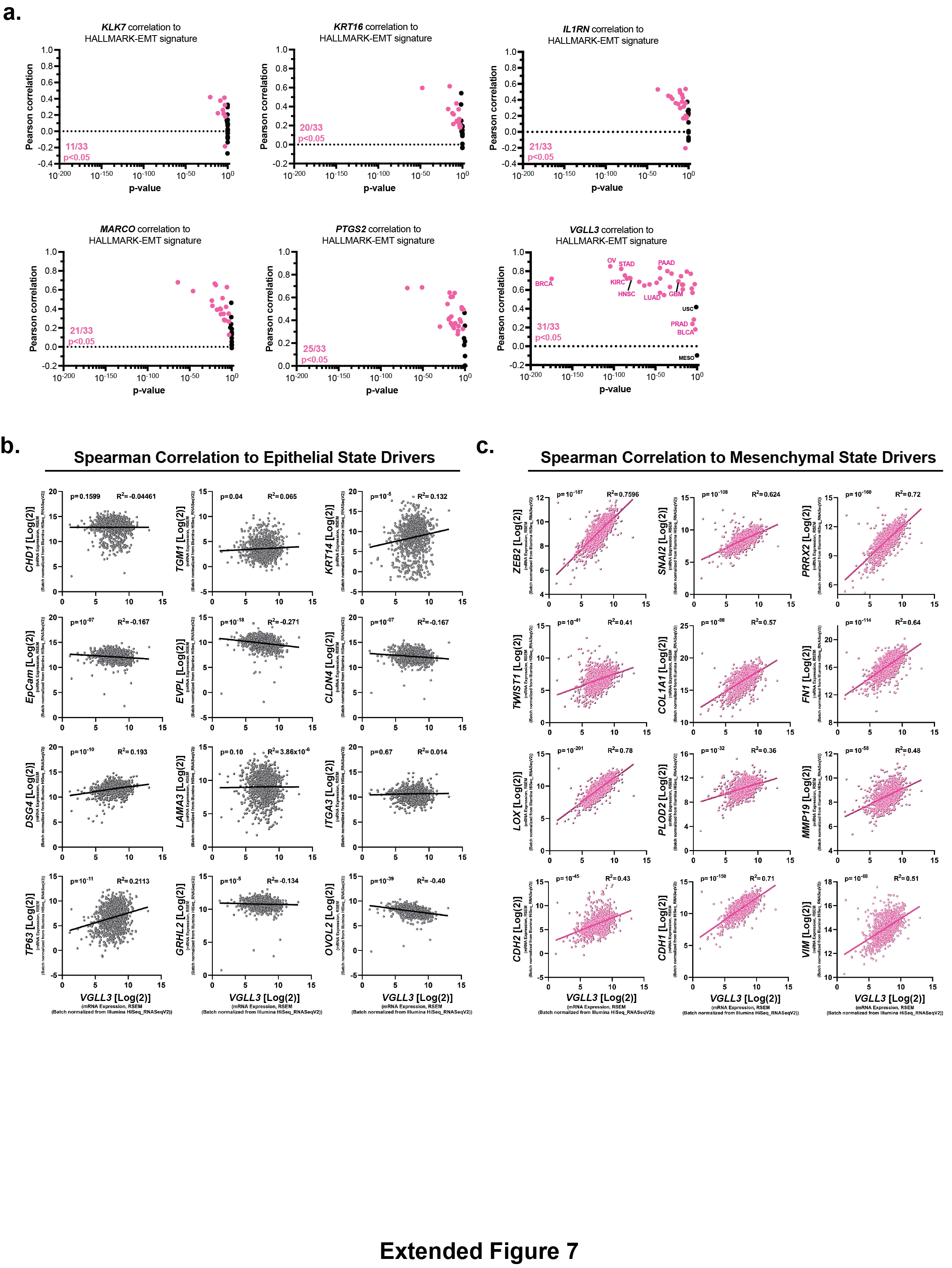

### Extended Figure 8

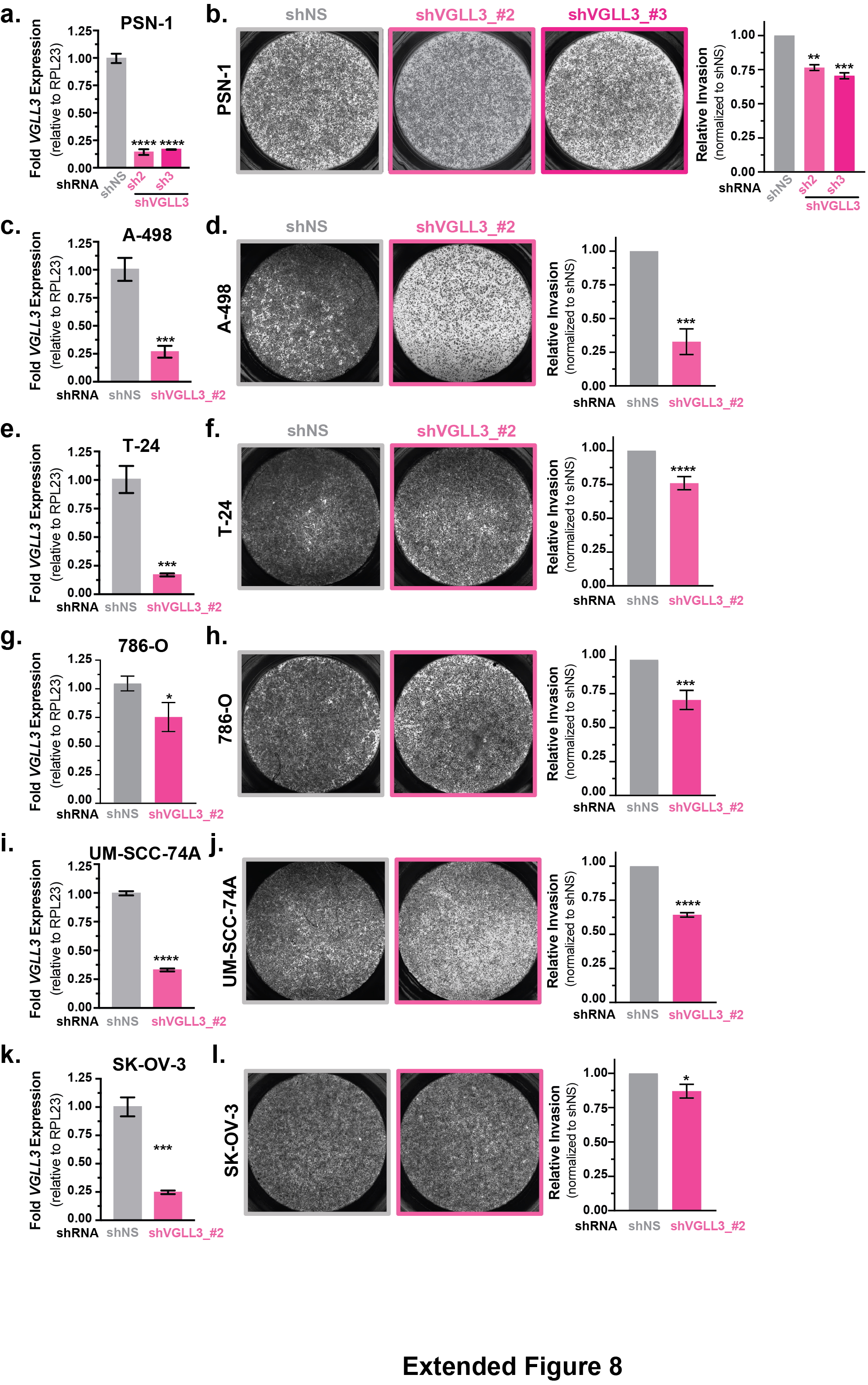

### Extended Figure 9

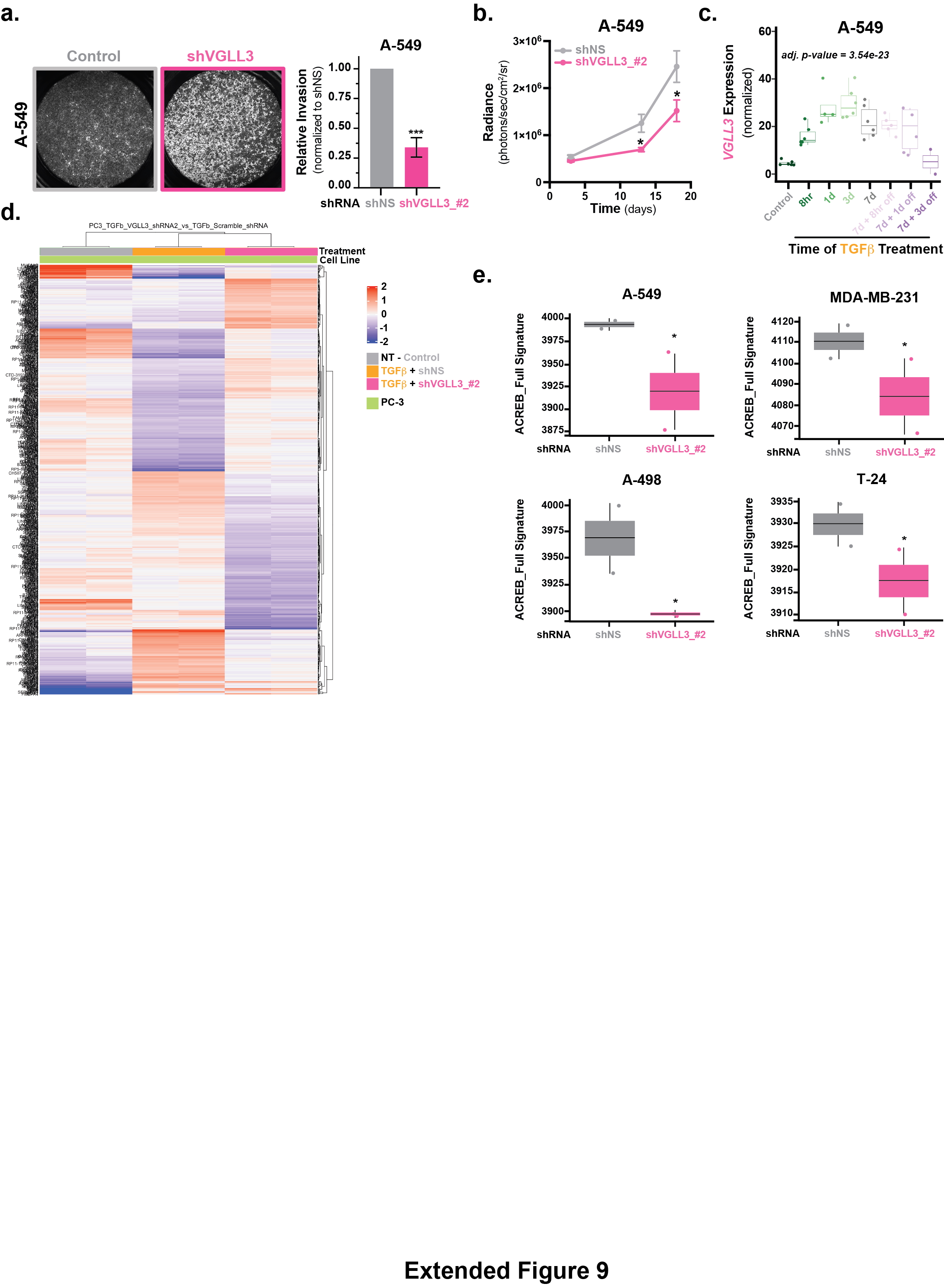

### Extended Figure 10

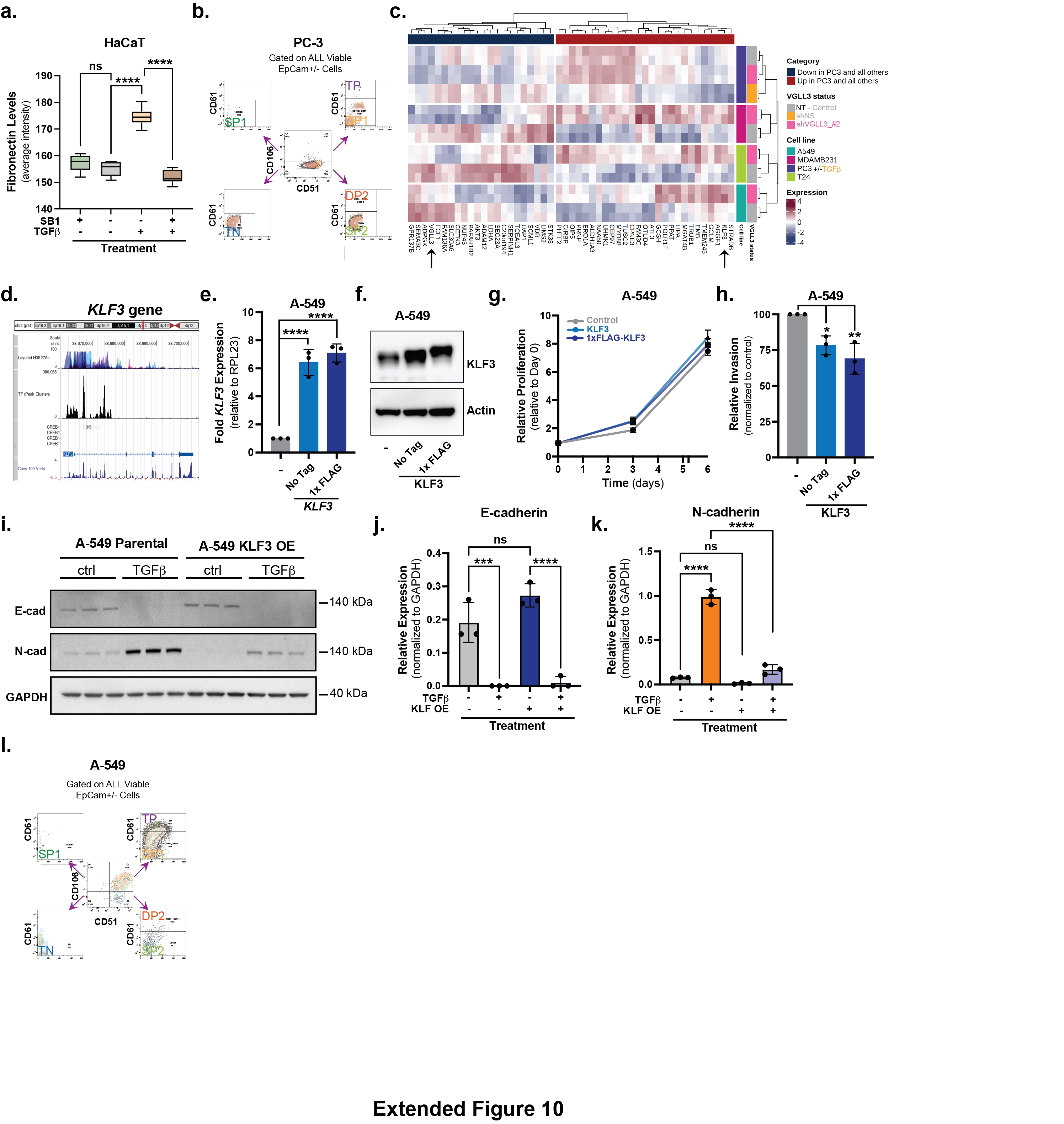
